# Elevational constraints on flight efficiency shape global gradients in avian wing morphology

**DOI:** 10.1101/2024.07.12.603304

**Authors:** Jingyi Yang, Chenyue Yang, Hung-wei Lin, Alexander C. Lees, Joseph A. Tobias

## Abstract

Wings with elongated shape or larger surface area are associated with increased flight efficiency and dispersal ability in a wide range of animals from insects to birds ^1–4^. Inter- and intraspecific variation in these attributes of wing shape is determined by a range of factors – including foraging ecology, migration and climatic seasonality ^5–8^ – all of which may drive latitudinal gradients in wing morphology ^9,10^. A separate hypothesis predicts that wing shape should also follow an elevational gradient because air density and oxygen supply decline with altitude ^11^, altering the aerodynamics of flight, and driving the evolution of more efficient wings in high-elevation species to compensate for reduced lift ^12,13^. However, previous analyses have found only mixed support for the ‘thin-air’ hypothesis ^14–18^, and we currently lack a global synthesis of elevational gradients in wing design for any taxonomic group. In this study, we use phylogenetic comparative models to explore elevational effects on wing morphology in 9986 bird species, while accounting for multiple climatic and ecological attributes, including latitude, temperature seasonality, body mass, aerial lifestyle and migration. We found that relative wing elongation (hand-wing index) and wing area increase with elevation, particularly in the upper montane zone (>4 km above sea level). These results confirm a pervasive elevational gradient in avian wing morphology, highlighting the role of aerodynamic constraints as key mechanisms shaping global patterns of trait evolution in flying animals.

## Results and Discussion

Morphological adaptations linked to dispersal mediate numerous fundamental processes including speciation ^19^, community assembly ^20^ and population-level responses to environmental change ^21–23^. The central role of dispersal in many aspects of ecology and evolution has led to increased interest in the mechanisms driving variation in dispersal ability, and the extent to which they can explain broad geographic patterns in organismal phenotype ^24,25^. The most prominent of these patterns is a latitudinal trend for increased dispersal ability, driven largely by global climatic gradients and their effect on species ecology ^9,26^. In flying animals, this trend is reflected in the increased expression of morphological flight adaptations from the equator to the poles ^9,27^, a pattern that has also been proposed for elevational gradients, with flight efficiency increasing from lowlands to mountaintops ^16^. However, this elevational trend in dispersal-related adaptation has received relatively little attention, and potentially arises from a different mechanism linking air density and the physics of flight ^12,28,29^.

Average air density decreases almost linearly from sea level to 10 km altitude ^11^, gradually increasing constraints on flight efficiency for aircraft and animals alike ^30,31^. In animals that rely on flight to forage or move across the landscape, this means that wing morphology adapted to lowland conditions may provide insufficient lift at higher elevations ^12,28^, thus driving wing-shape evolution to improve flight efficiency ^5,15,16,18^. Specifically, the ‘thin-air’ hypothesis predicts that wings should increase in elongation or area towards mountaintops ^5,16^ because longer and larger wings can improve energy efficiency or generate increased lift, respectively ^2,3,28,32^ (Figure 1). Despite the clear connection with well-established aerodynamic theory, current evidence for air density as a general mechanism driving wing evolution in animals is mixed and inconclusive ^17,33,34^.

**Figure 1.**
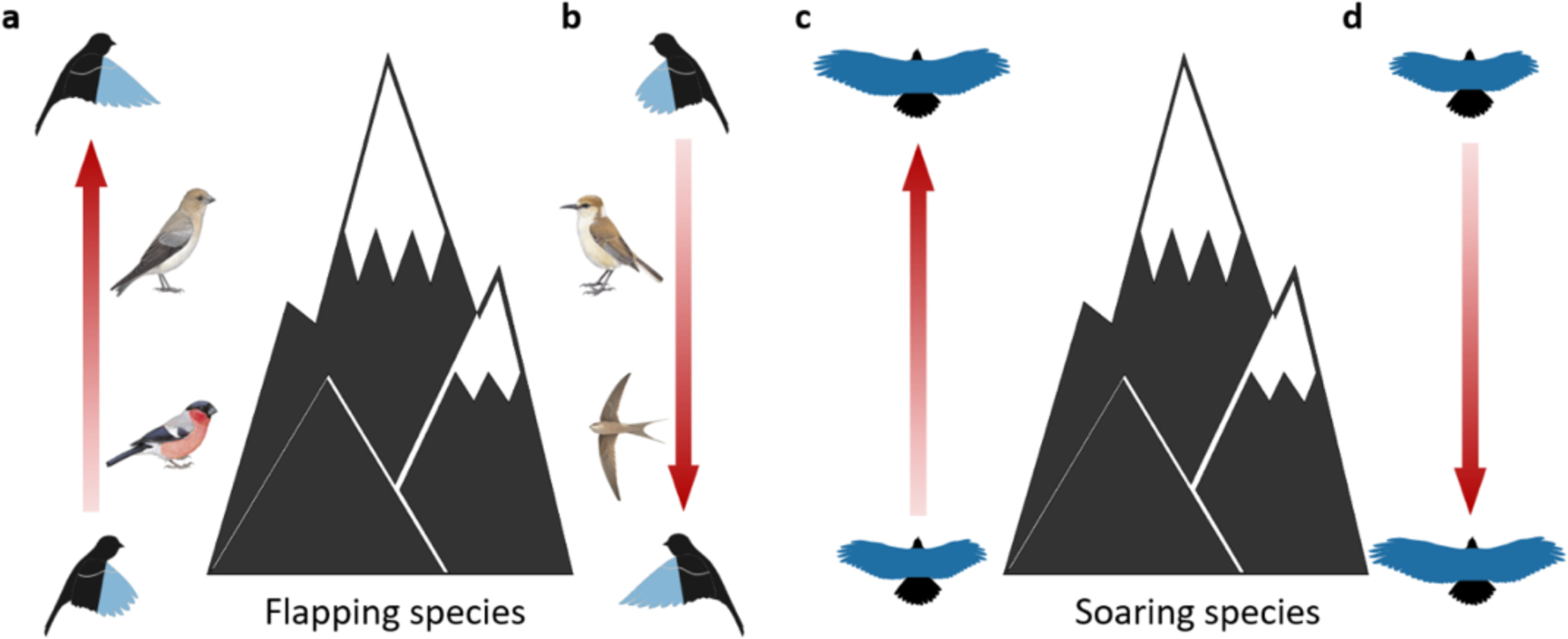
Potential direction and drivers of elevational gradients in avian wing morphology. (a) Air density declines with elevation from 0–10 km above sea level. Given that high flight efficiency is associated with more elongated wings, flapping species living at higher elevations (e.g., Sillem’s Rosefinch *Carpodacus sillemi*) may evolve longer and more pointed wings than lowland relatives (e.g., Eurasian Bullfinch *Pyrrhula pyrrhula*) to provide greater lift or higher flight efficiency in lower- density air. (b) Alternatively, gradients in flight adaptations can reflect foraging ecology and lifestyle, potentially reversing any elevational trend related to air density. For example, high mountains are colder with more open habitats and fewer flying insects, thus a higher proportion of montane species may be terrestrial (e.g., Ground Tit *Pseudopodoces humilis*), whereas lowlands are warmer with more flying insects, increasing the diversity of aerial insectivores (e.g., African Palm Swift *Cypsiurus parvus*). (c) Similarly, in the case of soaring species, larger or more elongated wings may evolve at higher elevations to compensate for lower air density and maintain flight efficiency. (d) Alternatively, the wing area gradient may be reversed because higher temperatures, and associated thermal uplifts, increase the size and diversity of soaring species at lower elevations. Red arrows show opposing elevational gradients – between lowlands and mountaintops – of wing length, hand- wing index (HWI) and hand-wing area (HWA). Images of exemplar bird species are reproduced with permission from Birds of the World (Cornell University).

The tendency of flying animals to have larger wings at higher elevations was recognised almost a century ago, although this pattern was often ascribed to a thermoregulatory mechanism, with wing-size differences emerging as a correlate of larger body size, in accordance with Bergmann’s rule ^35,36^. Early studies also speculated that lower air density could reinforce this pattern ^5,37^. For example, based on anecdotal observations of museum specimens, Erwin Stresemann ^38^ proposed that wings of high elevation bird species were longer and more pointed to compensate for the reduced ‘carrying capacity’ of the air. Recent quantitative analyses have tested variants of this hypothesis in a range of taxa, with some showing evidence of increased wing size or elongation at higher elevations for particular bird species ^15,39–41^, and other studies finding no such relationship ^42–44^. Similarly, in some groups of insects, wing size appears to increase in relation to body size at high elevations ^18^ whereas other studies show the opposite trend of wing reduction ^45,46^.

These opposing trends may reflect variation among species in the effects of ecological gradients linked to climate or food supply. For example, some insect species are thought to become less aerial and therefore shorter-winged at high elevations because lower temperatures and stronger winds increase the cost and risk of flight ^45,46^. In birds, too, the proportion of aerial-foraging species may decline at higher elevations, either because cooler temperatures reduce the availability of airborne insect prey ^47^, or limit the production of thermal updraughts used by soaring species ^48^.

Differences in species ecology may therefore reverse the pattern of wing elongation or enlargement at higher elevations in some contexts (Figure 1), complicating or biasing elevational trends in phenotype. In the most extensive study to date, Youngflesh et al. ^14^ found that relative wing length increased with elevation in a sample of 105 North American bird species (mainly passerines), although several alternative hypotheses were not tested, and it remains unclear whether this pattern holds true across broader geographic and phylogenetic scales.

To answer this question, we compiled a dataset of elevational ranges for all extant birds (n = 9986 species) to assess the direction and drivers of elevational gradients in wing morphology at a global scale. Specifically, we use phylogenetic models to test whether the maximum elevation at which bird species occur predicts their relative wing elongation and wing area. Birds offer an ideal opportunity to test the thin-air hypothesis as they populate the entire elevational range habitable to terrestrial vertebrates from sea level to at least 8.3 km altitude. In addition, the aerodynamic properties of their flight can be quantified using published morphological trait data ^49^ to calculate relevant metrics, including hand-wing index (HWI) and hand-wing area (HWA) ^1,9,50,51^. HWI – a measure of wing pointedness or elongation correlated with wing-aspect ratio ^52^ – is calculated from the lengths of the wing chord and the first secondary feather (Figure S1) which indicate wing length and wing width, respectively ^9,32,52^. Using the same variables, we also adapted a method proposed by Wright et al. ^50^ to compile a dataset of HWA for all birds. Although HWA quantifies the area of the outer wing only ^53^, we show that it is highly correlated with the aeronautical wing area – the total area of the underside of both wings plus the intervening underside of the body (see Methods; Figure S1-S2). Therefore, increased HWI is associated with greater energetic efficiency of flight while increased HWA indicates greater lift ^1,9,15,28^.

The influence of air density on HWI and HWA may vary according to flight style ^28,54^ (Figure 1), so we divide birds into groups that primarily use flapping flight (n = 9563 species) and soaring flight (n = 378 species). In combination, these datasets allow us to assess the impact of elevation on wing morphology in the context of multiple factors known to shape wing evolution, including body mass ^55^, flight mode and aerial lifestyle ^1,12,54,55^, migration ^6,26^, and trophic niche ^55,56^, as well as environmental variables such as latitude ^9^, temperature seasonality ^5,9^, and habitat openness ^5^ (see Methods). Thus, although it is not possible to directly test the impact of thinner air on flight adaptation at this scale, our analyses are able to disentangle the likely contribution of air density from other critical factors, many of which vary with elevation.

### Elevational gradients in avian wing morphology

Globally, we found that soaring birds are generally restricted to lower elevations and have more pointed and larger wings relative to body mass, compared with flapping species (Figure 2a,b). In both cases, soaring and flapping species show an overall increase in relative HWI and HWA as elevation increases, especially >4 km above sea level. However, there are prominent differences at low elevations, with soaring species showing a dramatic decrease in HWI and increase in HWA up to around 2 km above sea level (Figure 2a,b). This pattern is primarily driven by a subset of seabirds – including albatrosses and petrels – with dynamic soaring flight associated with extremely long and narrow wings ^57^. Unlike thermal soaring, dynamic soaring relies on strong winds and is largely restricted to marine environments at the lowest elevations (see Methods). When seabirds (n = 237 species) are removed, soaring and flapping landbirds show similar gradual increases in HWI and HWA with elevation, consistent with predictions of the thin-air hypothesis (Figure 2a,b). This evidence is further corroborated by a significant positive elevational gradient in average HWI and HWA when all species (including seabirds) are partitioned into elevational bands based on their distribution (Figure 2c,d).

**Figure 2.**
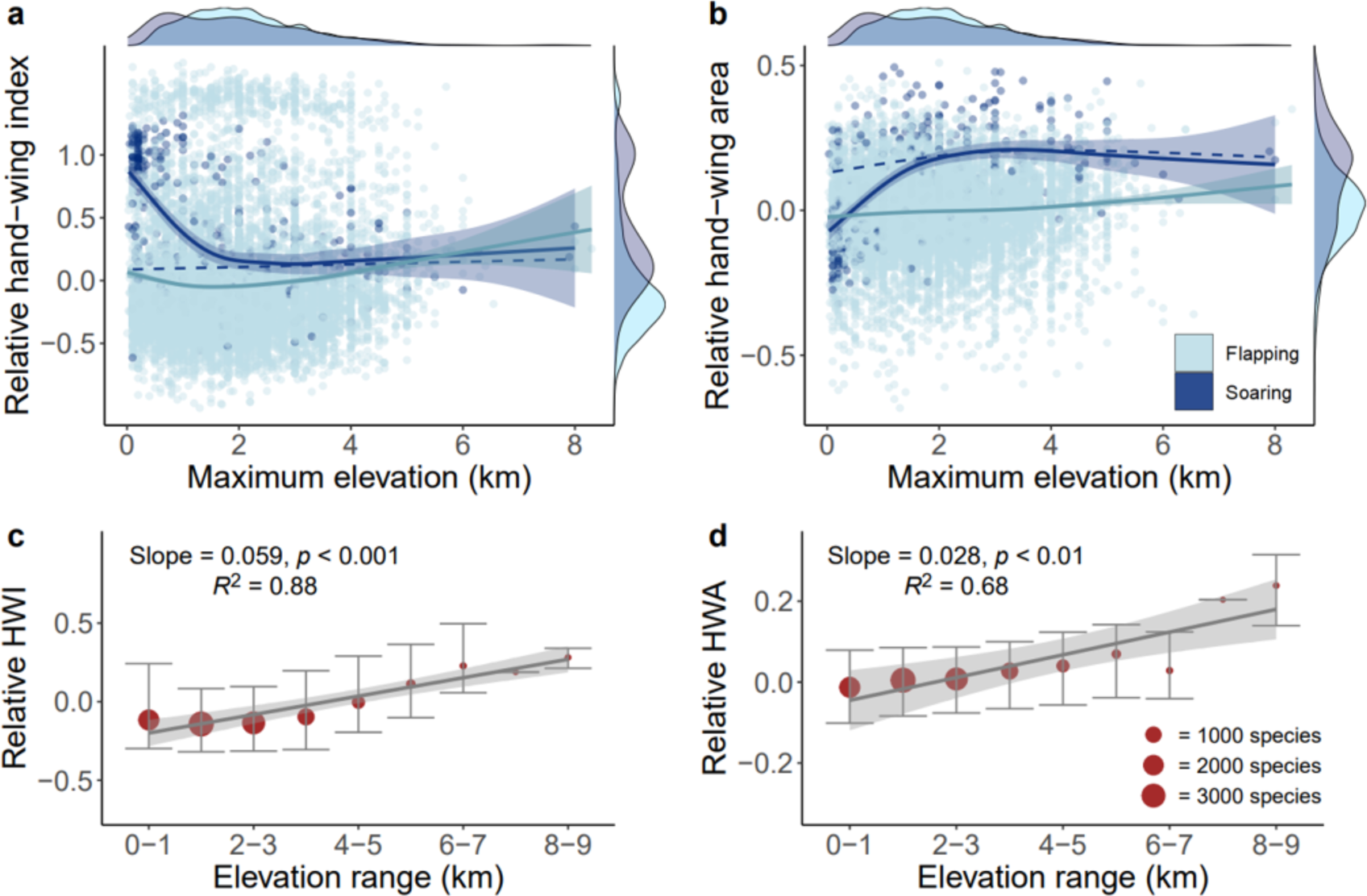
Relationship between wing morphology and elevation in birds. (a) In 9563 species with flapping flight, hand-wing Index (HWI) increases with maximum elevation (i.e. the highest elevation reported for each species). In a smaller sample (n = 378) of soaring species, HWI initially declines sharply with elevation under 2 km, and then increases at higher elevations. However, high HWI in low-elevation species is largely driven by narrow-winged seabirds, in which soaring is often wind- assisted and therefore less constrained by air density. After removing seabirds, soaring species show a more gradual increase in HWI with elevation (dotted line). (b) Hand-wing area (HWA) of flapping and soaring species also increases with elevation. When seabirds are removed (dotted line), HWA of terrestrial soaring species increases gradually up to 4 km, then levels off at the highest elevations. (a- b) Density plots adjacent to panels show the overall distribution of flapping and soaring species across axes of elevation and wing morphology. (c-d) Both relative HWI and HWA increase with elevation above sea level when flapping and soaring birds are grouped into 1 km-wide elevational bands based on their maximum elevation. Central dots show median wing measurements for each elevational band; whiskers show first and third quartiles; dot size represents the sample size within each elevational band (1–3382 species). Statistics shown are from a linear regression (grey line) between median HWI (c) or HWA (d) and max elevation; shaded area shows 95% CI. All analyses use relative wing measurements (HWI or HWA corrected for body size) calculated as the residuals from linear models between standardised wing measurements and body mass. HWA and body mass were log-transformed before standardisation.

The global-scale correlation between elevation and wing shape may be at least partly explained by factors other than air density, such as climate and species ecology. To tease apart the contribution of potential alternative mechanisms, we ran a set of phylogenetic generalised least square (PGLS) models to test the effect of elevation while controlling for eight major determinants of wing morphology (latitude, temperature seasonality, habitat openness, body mass, flight mode, aerial lifestyle, migration and diet). Even when accounting for the combined contribution of all these factors, we found that elevation remained significantly positively correlated with both HWI (95% CI = [0.000, 0.013], *P_t_* < 0.05) and HWA (95% CI = [0.017, 0.024], *P_t_* < 0.001; Figure 3, Table S1-S2) in models containing all volant bird species (n = 9787). To assess whether these results were driven by special cases, we removed two problematic subsets of species, namely seabirds, with their wind-assisted flight style, and long-distance migrants, which potentially occur outside their optimal range as vagrants or fly at extreme altitude during their migratory journeys ^58,59^ (see Methods). When we restricted our sampling to non-migratory landbirds (n = 8703 species), the effect of elevation remained similar for wing area (Table S2) and increased for HWI (95% CI = [0.002, 0.015], *P_t_* < 0.05; Table S1), with the overall findings supporting our all-species models (Figure 3).

**Figure 3.**
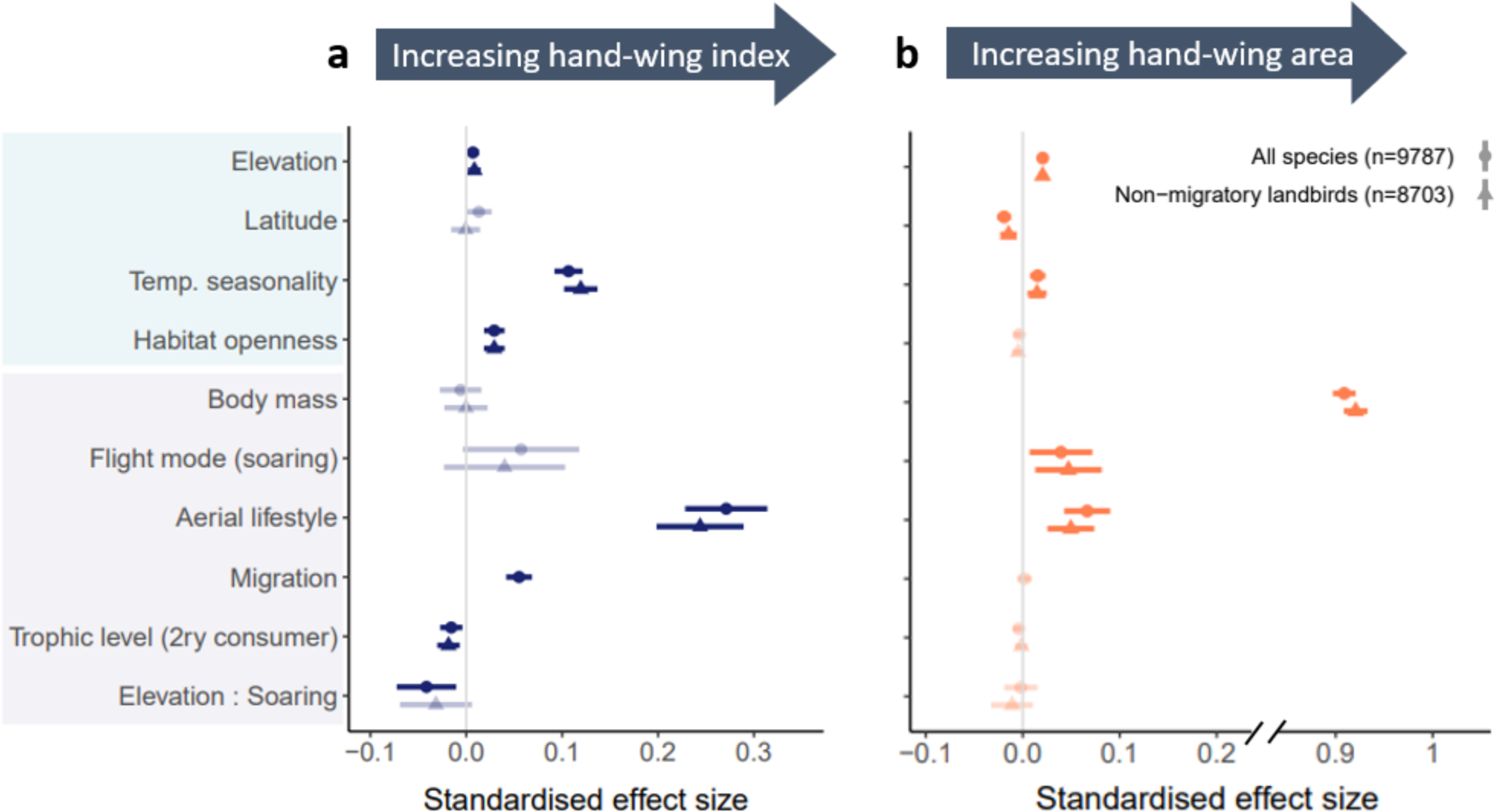
Phylogenetic models reveal that elevation predicts avian wing morphology. Forest plots show the parameter estimates of phylogenetic models testing effects of environmental factors (blue box) and species traits (mauve box) on hand-wing index (a) and hand-wing area (b). Central points show the mean; error bars show 95% CI. Results were generated using 100 randomly selected trees (obtained from BirdTree.org) and averaged via Rubin’s rules. Positive values indicate a positive correlation between the predictor and the corresponding wing metric. Seabirds and migratory species may be under different selection pressures because of wind-assisted flight and high-elevation migration, respectively, so we repeated models with those categories removed (i.e. restricting to non-migratory landbirds). In each panel, we show results based on a sample of all volant birds (n = 9787 species; dots) and the subset of non-migratory landbirds (n = 8703 species; triangles). Significant effects are inferred when estimated 95% CI do not span zero (highlighted in a darker shade).

Our models also confirm that changes in wing elongation and wing area with elevation are not driven by increases in body size, in contrast to previous claims ^35,36^. Indeed, when we removed body mass from model predictors to examine morphological changes without size-correction, we detected stronger positive effects of elevation on most wing metrics (Table S3-S6). This implies that body mass increases with elevation, in accordance with Bergmann’s rule, but more slowly than the increases in wing size. Thus, our findings confirm that birds living at higher elevation have longer and larger wings, on both absolute and relative scales, supporting the view that montane environments have general and consistent effects on the evolution of flight adaptations.

In soaring species (n = 378), the direction of morphological responses to elevation was less clear. Overall, their HWI seemingly decreased with maximum elevation (95% CI = [-0.065, -0.004], *P_t_* < 0.05) while HWA increased (95% CI = [0.002, 0.035], *P_t_* < 0.05; Tables S1-S2). However, the results of these models are again likely driven by marine species. When we removed all seabirds from the sample, both HWI and HWA were no longer associated with elevation (95% CI = [-0.060, 0.014] and [- 0.012, 0.031], respectively; Tables S1-S2). This removal of significant effects may simply reflect a much smaller sample size. Alternatively, divergent responses between flapping and soaring birds may be caused by their fundamentally different sources of lift during flight. Soaring species often generate lift from thermal convection currents, which are largely determined by temperature rather than air density ^48^. In addition, a weaker link between air density and wing morphology in soaring species at high elevations may arise because they rely on upwash – i.e., winds pushed upward by topographical features including slopes and cliffs ^104^ – or because they use other adaptations to improve lift ^105^, including respiratory air sacs ^60^.

### A steepening of morphological gradients towards mountaintops

Although our global models reveal overall trends in wing morphology across the entire elevational gradient, they provide only limited insight into how these trends vary with elevation. To explore the finer-scale and potentially non-linear relationship between wing morphology and elevation (Figure 2), we used a ‘sliding window’ approach to divide our sample into a series of subsets based on species’ maximum elevation (0-3 km, 1-4 km, and so on), then repeated PGLS models within each elevational band, accounting for variation in multiple climatic and ecological factors (see Methods). These ‘sliding-window’ models revealed that positive correlations between elevation and nearly all wing metrics gradually strengthened as elevation increased, with particularly strong effects over 4 km above sea level (Figure 4). For example, the effect of maximum elevation on HWI is weak or non-significant in the lowest five elevational bands, and jumps to a strong effect (0.252; 95% CI = [0.061, 0.443], *P_t_* < 0.05) in the highest elevational band. The findings are also robust to the removal of seabirds and migratory species (Figure S3), as well as using alternative methods of dividing the sample (see Methods; Figure S4). This steepening gradient suggests that avian wing morphology is more sensitive to elevational change at the highest elevations reached by birds, matching anecdotal observations of very long-winged species in the highest mountain ranges, e.g. *Muscisaxicola* ground- tyrants in the Andes, *Leucosticte* rosy-finches in the Rockies, *Carpodacus* rosefinches in the Himalayas (Figure 1). These cases are most likely evolutionary responses to reduced flight performance at high elevations, which in turn appears to be linked to reduced air density, rather than, for example, reduced oxygen availability ^31^. Nonetheless, oxygen deficiency may be a contributory factor increasing the negative impact of lower air density towards mountaintops ^16,18^, thereby accentuating selection for optimal flight efficiency.

**Figure 4.**
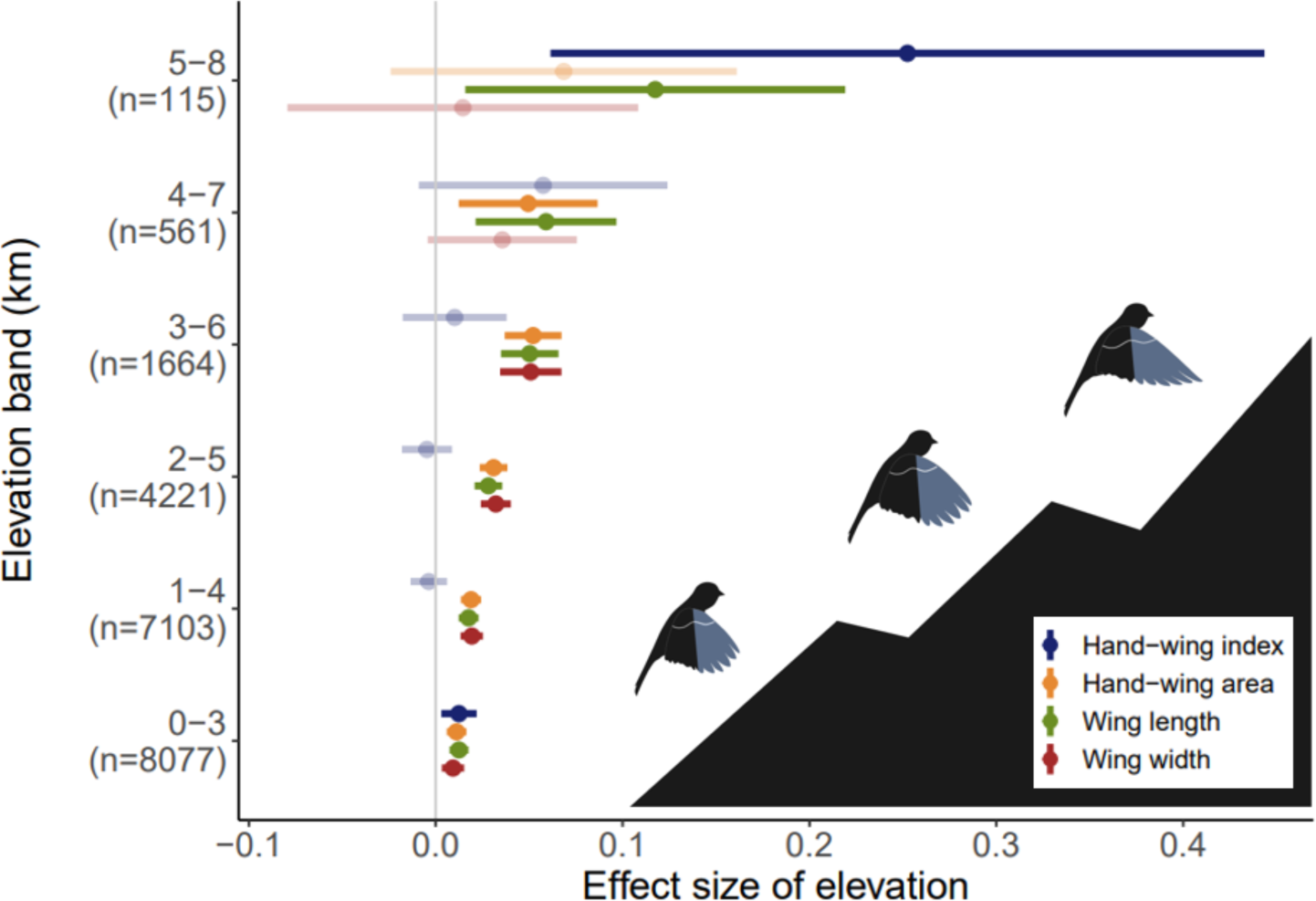
Gradients in avian wing morphology are accentuated at the highest elevations. Forest plot shows the parameter estimates of maximum elevation on wing morphology, based on separate phylogenetic models for each 3-km wide elevational band (starting with 0-3 km and ending with 5-8 km above sea level). The sample sizes within elevational bands refer to flapping species only, because the models included an interaction term (Elevation : Flight mode) such that the main effect of elevation was estimated with flight mode at the reference level. Central points show the mean; error bars show 95% CI. Results were generated using 100 randomly selected trees and averaged via Rubin’s rules. Significant effects are inferred when estimated 95% CI do not span zero (highlighted in a darker shade). The findings were robust to the removal of seabirds and migratory species (Figure S3) and the use of alternative bandwidths (2km and 4 km; Figure S4). Silhouettes illustrate tendency of wings to become larger and more elongated with increasing elevation.

To obtain further insights into adaptive mechanisms, we decomposed our flight efficiency metrics into their underlying variables: wing length and wing width (Figure S1a). In both cases, the global PGLS models showed a significant positive relationship with maximum elevation (Figure S5). In other words, increasing elevation is not simply associated with wing elongation but also with an increase in wing width, helping to explain why the effects of elevation are not restricted to HWI (an index of elongation) and more strongly predict the positive gradient in HWA (which reflects a combination of wing length and width; Table S1-S2). A closer inspection of elevational gradients in these traits reveals that wing length and wing width (i.e. secondary 1) both increase at similar rates for the first 4-5 km above sea level, after which the slope in wing width flattens while the slope in wing length increases (Figure 4). This helps to explain the abrupt increase in HWI at extremely high elevations.

## Implications and applications

Indices of flight efficiency (e.g. HWI) are often assumed to reflect dispersal ability or dispersal limitation in birds ^9,22,50,51^. While this assumption is likely to be robust in macroecological studies ^1^, our findings suggest a potential decoupling of wing shape and dispersal ability at high elevations. That is, the prominent increase in HWI or wing area at mountaintops may reflect a compensatory adaptation that maintains the same levels of aerodynamic capacity without necessarily translating into increased flight efficiency or dispersal ability. Variation in wing shape should therefore be used with caution as a proxy for these traits, particularly when sampling across broad elevational gradients. In particular, analyses using avian HWI as an index of flight efficiency should consider accounting for air-density effects using global datasets of avian elevational ranges presented in this study (Data S1).

Our study may also have implications for understanding and predicting responses of montane biodiversity to climate change. Global warming has long been known to drive upslope movement of animals ^61^, with dispersal ability being a critical factor determining the rate and extent of elevational shifts to cooler montane refugia ^62^. A growing number of studies have explored the behavioural and physiological challenges facing lowland species tracking their climate niches upslope ^63,64^, particularly in the tropics ^65,66^. Previous analyses have highlighted a range of obstacles to life at high elevations, including competitive interactions ^67,68^, impaired breeding and metabolic inefficiency ^69–71^. Our findings add a further dimension to these challenges, suggesting that aerodynamic factors will increase flight costs for species adapted to lowlands, potentially capping elevational range shifts. Similar constraints may help to explain the counter-intuitive findings of a previous study showing that flying insects are slower, rather than faster, to shift upslope due to the higher aerobic demands of flight, exacerbated by lower air density ^72^.

## Limitations and future directions

A number of factors should be borne in mind when interpreting our results. For example, the behaviour of many bird species is poorly known, particularly in the tropics, increasing uncertainty around our classification of flight behaviour, especially aerial lifestyle. In addition, we use the maximum elevation for each species, which has many advantages (see Methods) but may be more sensitive to sampling effort, and potentially overemphasises extreme conditions that are not experienced by the majority of individuals. To explore these issues, we scored data uncertainty using standard techniques ^73^, and calculated mean elevation for each species (see Methods). When we re-ran all models restricting the sample to high-certainty data (n = 6864 species) or replacing maximum elevation with mean elevation (n = 9787 species), the results remained largely unchanged (Table S7- S10, Figure S6). The relationship between elevation and HWI was slightly weaker using these alternative methods, suggesting that HWI (wing elongation) constrains the maximum elevation reached by birds, and is less strongly associated with average elevation. The correlation between elevation and all other wing metrics (wing area, length and width) remained the same (Table S7-S10), supporting our main conclusions.

After accounting for a range of contributory factors, the absolute effect of elevation on wing morphology in our models was relatively small (Figure 3). While this may seem surprising given that air density declines by >65% from sea level to 10 km altitude ^11^, the effect on wing shape may be small because other flight adaptations could evolve in response to low air density and low oxygen levels. For example, buoyancy can be aided by concealed air sacs in soaring birds ^60^, while all flying animals can theoretically improve flight efficiency in thin air by adjusting the ‘angle of attack’ in their wing movements ^74,104^, altering flight speed and trajectory ^12,75^, or morphing wing shape during flight to achieve higher manoeuvrability ^76,104^. Such behavioural plasticity can potentially reduce the physical and energetic costs of low air density, thereby dampening selection on wing morphology ^74^.

The relationship between air density and wing evolution could also be complicated by a suite of other physical properties known to influence flight performance, including air temperature, humidity and wind speed ^16^. While average air density decreases almost linearly across the altitudes relevant to avian flight, air density itself is influenced by temperature and humidity, and thus may partially decouple from altitude, giving rise to the concept of ‘density altitude’ in aeronautics ^77^. In addition, wind speed generally increases with altitude ^16^, potentially compensating for the decline in air density because lift is proportional to wind speed ^12^. To date, most aerodynamic models are designed to describe stable or gliding flightpaths, while living organisms need to optimise flight during various stages (e.g. take-off and landing ^78^) as well as complex atmospheric conditions ^79,80^.

Thus, a more thorough understanding of the impact of air density on animal flight adaptations requires the development of more refined aerodynamic models, along with a consideration of multiple alternative pathways by which birds can accentuate the support and thrust provided by the wings in flight ^104^. Finally, experimental manipulations of atmospheric properties and wing morphology in a sample of different organisms may provide a more rigorous test of the thin-air hypothesis and associated ideas.

## Conclusions

Previous research has confirmed that a latitudinal gradient in avian wing morphology is driven largely by ecological adaptations to climatic seasonality at higher latitudes, including accentuated dispersal ability and migration as strategies to cope with seasonal fluctuations in resources ^6,9,26^. Our results reveal that flight adaptations also follow a consistent elevational gradient at a global scale, with both wing elongation and area increasing towards mountaintops. This pattern persists even when we account for a wide range of climatic and ecological factors predicted to shape avian wing morphology. Our findings therefore provide compelling support for the thin-air hypothesis, particularly given the steep gradient in wing length detected at the highest elevations reached by birds, where a combination of lower air density and reduced atmospheric oxygen may exert strong selection for greater lift or flight efficiency. More generally, our findings highlight the potential role of aerodynamic constraints as a general mechanism driving wing evolution in flying animals and limiting their capacity to undergo rapid range shifts to higher elevations.

## Acknowledgements

We are grateful to Paul French for the initial inspiration to investigate this question, and to Rob Barber and Tom Weeks for help with data curation and analyses. Data collection was supported by the Silwood Park master’s programme. Publication costs were funded through the Imperial Open Access Fund.

## Author contributions

J.A.T. and A.C.L conceived the project; J.Y., C.Y. and H.L. collected data; J.Y. conducted formal analyses and visualisation; J.Y. and J.A.T drafted the original manuscript; all authors contributed to manuscript reviewing and editing.

## RESOURCE AVAILABILITY

### Lead contact

Further information and requests for resources should be directed to and will be fulfilled by the lead contact, Jingyi Yang.

### Materials availability

This study did not generate new unique reagents.

### Data and code availability

All data and code for reproducing this study are available at xxxx [url to the data & code repository will be added after acceptance].

## EXPERIMENTAL MODEL AND SUBJECT DETAILS

### Morphological trait data

To quantify interspecific variation in wing morphology related to flight ability, we used two metrics: hand-wing index (HWI) and hand-wing area (HWA). We extracted HWI from global datasets ^9,49^ that calculated the index from two wing measurements – wing length (WL) and the first secondary length (SL) – as (WL-SL) / WL x 100 following ref. ^19^ (Figure S1). HWI describes the pointedness or elongation of the wing, as opposed to wing length; e.g. hummingbirds (Trochilidae) have relatively short wings with high HWI whereas trumpeters (Psophidae) have relatively long wings with low HWI. Unlike wing length, HWI therefore provides an index of wing-aspect ratio ^52^ reflecting flight efficiency and dispersal ability ^1,9^. Species averages for HWI, WL and SL were obtained from the AVONET dataset ^49^ (n = 9986 species) wherein most estimates are based on samples of at least four museum specimens per species, or inferred from closely related species with similar morphology. Inference was required in only 45 species (0.5% of the sample) lacking measurements for one or more traits, and the identity of surrogate species is provided in AVONET ^49^.

No global datasets of avian wing area currently exist, so we compiled standardised estimates of hand-wing area for all species using WL and SL. Standard aerodynamic models ^28^ define wing area as the total area of the underside of both extended wings plus the underside of the intervening body (Figure S1). However, we were unable to replicate this measurement because it is not possible to open or extend the wings of museum specimens. In addition, we were unable to use the wing area calculation suggested by ref. ^53^ because wingspan measurements are not readily available for most bird species. Instead, as a proxy, we calculated the area of the ‘hand’ portion of both wings (hereafter termed hand-wing area, HWA) using the formula proposed in ref. ^50^: WL x SL x π/2 (Figure S1). To evaluate the validity of HWA as a proxy, we compiled published estimates of the total aeronautical wing area measured from live individuals or spread-winged specimens of 630 bird species. Even though HWA accounts for a relatively small portion of the total wing area (Figure S1), we found that HWA and total wing area are closely correlated (Pearson’s correlation = 0.99; Figure S2). This tight relationship supports the use of HWA as an index of total aeronautical wing area in our analyses and future studies. We present our full dataset of HWA for 9986 bird species in Data S1.

### Elevational distribution

To compile maximum and minimum elevation data for all extant bird species, we began by merging data from two large datasets ^82,83^. When these sources lacked data for a particular species, or provided estimates that differed by >0.3 km, we filled gaps and resolved conflicts using a third dataset ^81^ in conjunction with other published literature. Where multiple elevational ranges were given for one species in any source (e.g. for different subspecies or localities, such as ref. ^83^), we selected the extreme maximum and minimum values reported across the entire distribution of the species. To resolve disparities, we checked errors arising from inaccurate records or taxonomic mismatches by consulting online resources, including Birds of the World ^96^ and eBird citizen science data (www.ebird.org), before merging data from multiple sources and calculating the final elevation range for each species.

We applied several steps to overcome incompatibilities between sources of elevation data. First, data from ref. ^82^ are presented on a log scale, so we back-transformed data from that source and rounded to integers. Second, we omitted erroneous or uncertain estimates. For example, ref. ^83^ provided a maximum elevation of 5895 m for Preuss’s Weaver *Ploceus preussi*, a resident species restricted to lowland rainforests in West Africa. We revised the maximum elevational limit for this species downward to 1 km above sea level based on eBird localities (www.ebird.org). In migratory or wide-ranging species, we allowed a larger leeway for uncertainty because extralimital high- or low- elevation records are more probable. Third, we ensured that the seasonal context of elevational range data was compatible. Two core sources ^81,82^ provided elevation estimates from the total geographical range of each species, whereas a third source ^83^ provided estimates from the breeding range only. This mismatch has limited effect at the scale of our analysis because ∼80% of bird species are non-migratory ^49^, while many migratory species reach their highest elevation during the breeding season. When we compared samples of species for which both types of data were available, we found that elevation data based on the breeding range is strongly correlated with data from the year-round range (Pearson’s correlations for max elevation: 0.92, n = 7087 species; for min elevation: 0.94, n = 6102 species; Figure S7). Nonetheless, to minimise inconsistency, we re-checked all migratory species with elevation data based exclusively on ref. ^83^ to ensure their maximum and minimum elevations also included data from the non-breeding season. Finally, some data inconsistency between sources arose from taxonomic mismatches, which we resolved using resources including Birds of the World ^96^ and Avibase (https://avibase.bsc-eoc.org/). For instance, following an older taxonomy ^97^ ref. ^82^ presented a maximum elevation of 3100 m for Freckle-breasted Thornbird (*Phacellodomus striaticollis*), which has been recently split into two species with distinct elevational ranges (0-700 m for *P. striaticollis*; 1000-3100 m for *P. maculipectus*) ^96^. In such cases, we deleted erroneous data and ensured the final data agree with BirdTree ^86^ species delimitation used in our analyses.

After producing the updated and refined dataset, we selected the final species-level elevation range using two alternative methods. We gave priority to the most recent estimates ^81^ that incorporate the latest and most complete distributional range of the species (i.e., in the order of ref. ^81^ > ref. ^82^ > ref. ^83^ ; Method 1). This method is likely to produce more accurate estimates of minimum and maximum elevation for species, but potentially introduces subjectivity. We therefore resampled all species using an alternative approach (Method 2) which involved a simple extraction of the extreme minimum and maximum elevation reported for the species, with no preference for sources. We then calculated the mean elevation for each species as the midpoint between the selected maximum and minimum values, i.e., (max + min)/2, using both Method 1 and 2. The elevation data yielded from the two methods were highly correlated for both maximum elevation (Pearson’s correlation = 0.99) and mean elevation (Pearson’s correlation = 0.99; Figure S8). We therefore used our first dataset (Method 1) for all analyses. Final elevation data (Method 1 and 2) and data sources for all bird species (n = 9986) are listed in Data S1.

### Geographical and climatic data

Avian wing morphology varies with latitude ^9,98^, so the effect of elevation on flight adaptation can only be understood in the context of latitudinal effects. Geographical distribution of bird species was sourced from the expert-drawn range polygons provided by BirdLife International (version 2019.1) ^85^, restricted to resident and breeding ranges in areas where the species is coded as extant and either native or reintroduced. Given the potential taxonomic mismatches between BirdLife and BirdTree species, we adapted the BirdLife maps using the following methods: (1) when there was a one-to-one match between BirdLife and BirdTree species (n = 8949), we used BirdLife maps version 2019.1 for the corresponding BirdTree synonym; (2) when one BirdTree species is split into multiple BirdLife species we combined the corresponding BirdLife range polygons into a single polygon reflecting BirdTree taxonomy (n = 1929 BirdLife splits); (3) when multiple BirdTree species share one or multiple BirdLife species ranges we used an earlier version of BirdLife range maps (version 2.0, published in 2012) created prior to most taxonomic changes between BirdLife and BirdTree (n = 198 BirdLife lumps. See ref. ^49^ for more details).

Using the adapted maps, we extracted the centroid latitude for each BirdTree species (n = 9855 extant species with suitable maps) using R package ‘PBSmapping’ ^91^. To quantify the temperature seasonality experienced by each species, we first overlaid the same maps with the annual temperature seasonality (Bio4) raster from the CHELSA dataset (version 2.1) ^84^ at the resolution of 30 arcsecond. We then extracted the mean Bio4 value for each species by averaging grid cells that fell within the species’ range using a Behrmann equal area projection, disregarding cells with less than 50% overlap. Latitude and temperature seasonality data for 9836 bird species are provided in our final dataset (Data S1).

### Ecological trait data

Most previous studies describing elevational gradients in avian wing morphology (e.g. ref. ^14^) do not account for other factors known to influence wing shape evolution, including flight mode and aerial lifestyle. Aerodynamic constraints of air density may differ between different wing uses ^28,54^, so we also assigned each bird species to one of the flight mode categories (flightless, flapping, soaring).

Flight mode classification was based on published descriptions of flight behaviour and inspection of videos and photographs of flying birds, where available ^96^. Regardless of flight mode, the extent to which species use flight also plays a major role in shaping morphological flight adaptations ^1^, suggesting that elevational gradients in wing area and elongation may reflect variation in flight behaviour across different elevational ranges (Figure 1). To account for this variation, we adapted methods proposed by ref. ^1^ and classified each bird species according to an aerial lifestyle index, with scores ranging from 0-3 to reflect the increasing importance of flight in the daily routine of the species (0 = non-aerial; 1 = infrequent flier; 2 = moderate flier; 3 = frequent flier). Aerial lifestyle may have greater uncertainty for rare or poorly known species, so we also scored data certainty for our aerial lifestyle classifications, with certainty scores varying from A (high certainty) to C (low certainty), using methods described in ref. ^73^. We present this novel dataset of aerial lifestyle indices for all birds (n = 9986 species) accompanied with estimates of data certainty in Data S1.

Three other major factors influencing avian wing morphology – habitat, migration, and diet – are also rarely considered in studies of flight adaptation on elevation gradients. Species living in open habitats, such as grasslands, deserts and rocky landscapes, tend to be more mobile and dispersive than those inhabiting dense habitats, such as forests ^9^. Thus, we classified each species as living in dense or open habitats, and used this binary variable to represent habitat openness. Long-distance migration also strongly influences wing morphology, particularly HWI ^9^, so we included migration as a binary predictor, with all species classified as either non-migratory (sedentary species, partial or altitudinal migrants) or migratory (long-distance migrants). For species with both migratory and non- migratory populations, they were assigned to the category relevant to the largest population by geographical area. From a dietary perspective, species occupying different trophic levels may have different foraging strategies or home range sizes, driving variation in flight behaviour ^55,56^. We therefore included trophic level as a binary predictor consisting of primary consumers (herbivores and omnivores) and secondary consumers (carnivores and scavengers). Finally, raw measurements of wing morphology are correlated with body size, so we included body mass as an explanatory variable in models to assess the changes of wing elongation and area in relation to overall body size ^99^. Data on habitat openness, migratory behaviour, trophic level, and body mass were obtained or adapted from AVONET ^49^, with detailed descriptions of all variables provided in Data S1.

### Phylogenetic data

Global bird phylogenies were downloaded from BirdTree.org ^86^ using the Hackett topology ^100^. We used the full version containing all species (n=9993) to maximise our taxonomic coverage. 100 trees were then randomly selected out of the 10k trees and this subset was used in all subsequent analyses.

## QUANTIFICATION AND STATISTICAL ANALYSES

All statistical analyses and visualisation were implemented in R ^87^ version 4.2.2.

### Phylogenetic modelling

After removing flightless taxa, our sample contained 9941 extant bird species. To assess the relationship between elevation and wing morphology, while accounting for evolutionary relationships among these species, we used phylogenetic generalised least square (PGLS) models implemented with the R package ‘phylolm’ ^93^. Our main models test whether interspecific variation in a flight-related morphological trait (HWI or HWA) can be predicted by the maximum elevation reached by each species. We used maximum elevation as our main predictor because the air density gradient is likely to constrain flight performance most acutely at the uppermost elevations. The value of maximum elevation per species also suffers from less ambiguity than other metrics such as mean or median elevation, given the largely unknown population distribution of most species. Multivariate models were used to account for the effects of alternative explanatory variables, including elevation and eight additional environmental and ecological predictors (i.e., latitude, temperature seasonality, habitat openness, body mass, flight mode, aerial lifestyle, migration and trophic level). Detailed definitions of all variables used in this study are presented in Data S1. To standardise variable types, we combined categories 1 and 2 of aerial lifestyle index and used it as a binary variable. We also added an interaction term between elevation and flight mode because in flapping flight and soaring flight, lift is generated through different mechanisms at different energetic costs to the bird ^28,54^, and therefore flight mode may induce divergent aerodynamic effects at the same air density. To further explore the details of wing morphological changes, we also modelled the change of wing length and width using WL and SL as the response variable. To assess the absolute change of wing morphology with elevation, we repeated models with body mass removed from explanatory variables.

In each analysis, we accounted for phylogenetic uncertainty by running models 100 times using the randomly selected trees. We then averaged parameter estimates using Rubin’s rules as recommended by ref. ^101^, implemented with the R package ‘mice’ ^94^. To account for allometric scaling ^102^, we log transformed morphometric values, including body mass, wing area and other linear measurements (WL, SL). We scaled all continuous variables to a mean of zero and standard deviation of 0.5 to enable direct comparison between the effect sizes of continuous and binary variables ^103^.

Response variables were also scaled to enable comparison between wing morphological metrics. Collinearity between explanatory variables was low to moderate (variance inflation factor: 1.00– 3.82), with the highest correlation found between latitude and temperature seasonality. We also assessed model fit with conditional R^2^ values calculated using the R package ‘rr2’ which accounts for the hierarchical data structure ^95^. We report any results where the 95% confidence intervals do not overlap with zero as statistically significant.

To examine finer-scale and potentially non-linear responses of wing morphology to elevation as shown in Figure 2, we used a ‘sliding window’ approach where species were assigned to a series of 3-km wide elevational bands according to their maximum elevation. The lowest band starts at sea level (0 km), with each subsequent band starting 1 km higher than the previous one (i.e., the first band spans from 0-3 km, the second band from 1-4 km, and so on, up to the highest band of 5-8 km). Note that bands are overlapping so a species may be assigned to multiple bands. For example, a species reported up to a maximum elevation of 1 km would appear in the first and second bands.

Only one species - Alpine Chough (*Pyrrhocorax graculus*) - had a maximum elevation above 8 km so we included it in the highest band. We then ran independent models within each of the six elevational bands using the same set of predictors.

### Sensitivity analyses

To examine potential biases in our dataset, we ran sensitivity analyses with marine and migratory bird species removed. Marine species (seabirds) are potentially anomalous because many of them specialise in ‘dynamic soaring’, a flight style that generates lift from the gradient of wind speed ^57^. This contrasts with terrestrial soaring species, most of which use ‘thermal soaring’ to generate lift from thermal convection currents ^48^. Another problem relating to marine species (including non-soaring species) is that a key predictor, temperature seasonality, is primarily based on land surface temperature and therefore the data is less accurate for seabirds. We removed migratory species because their maximum elevation data may be erroneous for two main reasons. First, some migrants fly at extreme altitude during migration, but estimates are only available for a handful of species with sufficient tracking data ^58,59^. Second, elevational data for some migrants may be exaggerated by occasional sightings of vagrants observed at high elevation during migratory journeys. Therefore, we avoid these sources of noise and uncertainty by running an initial sensitivity analysis focused on non-migratory landbirds only (n = 8703 species).

To further test the robustness of our results, we ran two additional analyses, one excluding species with uncertain flight behaviour from models, and another using mean elevation instead of maximum elevation as the response variable. Since the likelihood of assigning incorrect aerial lifestyle is increased in poorly known species, we repeated models while only retaining species with the highest-certainty aerial lifestyle data (certainty score = A; n = 6864 species) to assess the effect of data uncertainty. In the other analysis, we replaced maximum elevation with mean elevation because maximum elevation data are more sensitive to sampling effort and sometimes reflect extreme or unusual observations, while mean elevation may reflect the more typical air density experienced by the species. In addition, when running the second sensitivity analysis for our ‘sliding- window’ models, we also re-assigned species to elevational bands based on their mean elevation instead of maximum elevation in congruence with the model structure (i.e. mean elevation as the response variable).

We also explored the effect of our methods used to divide elevational bands in our final analyses. The results reported in Figure 4 are based on a bandwidth of 3 km, which was selected as a compromise between precision and sample size - smaller bandwidths tend to under-sample the variation within the band; larger bandwidths provide less detail about variation across the elevational bands. To assess whether our results are robust to variation in bandwidth, we re-ran our models using bandwidths set to 2 km and 4 km.

## SUPPLEMENTAL INFORMATION

Supplemental information can be found online at xxxx [url provided after acceptance]

## Supplemental Information

**Figure S1.**
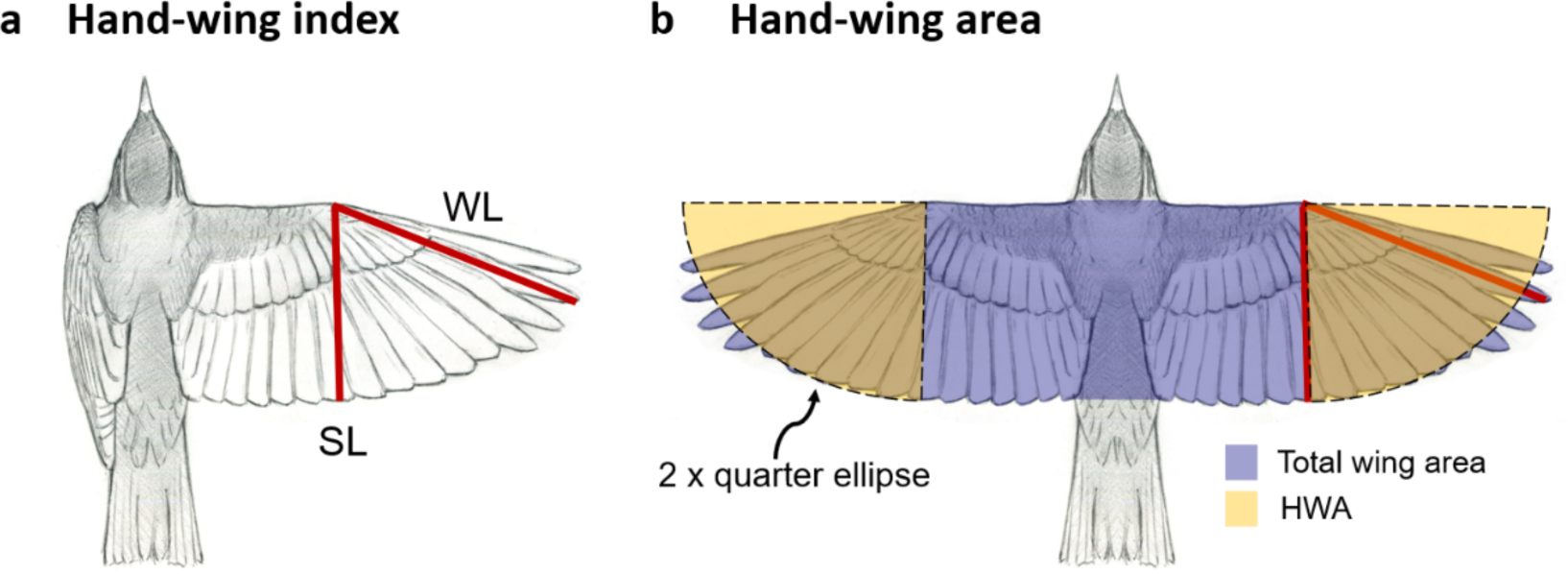
Wing morphological metrics used in this study. Wing length (WL): the length from the carpal joint (bend of the wing) to the tip of the longest primary. Secondary 1 length (SL): the length from the carpal joint to the tip of the first secondary feather, i.e. the outermost secondary adjacent to the innermost primary feather. (a) Hand-wing index (HWI) – an index of wing pointedness related to wing aspect ratio – is calculated as 100*(WL–SL)/WL. (b) Hand-wing area (HWA) is the estimated area of the outer wing, or hand-wing (yellow highlights). We adapted the formula proposed by ref. ^S1^ to calculate the area of both hand-wings: WL x SL x π/2. This formula assumes that the area of each of hand-wing can be approximated by a quarter of an ellipse, with two hand-wings therefore equating to half an ellipse (yellow highlights). HWI and HWA are widely used as metrics of flight ability in birds ^S1–3^. HWA is closely correlated with the total wing area defined by standard aerodynamic models ^S4^ (purple highlight; Figure S2).

**Figure S2.**
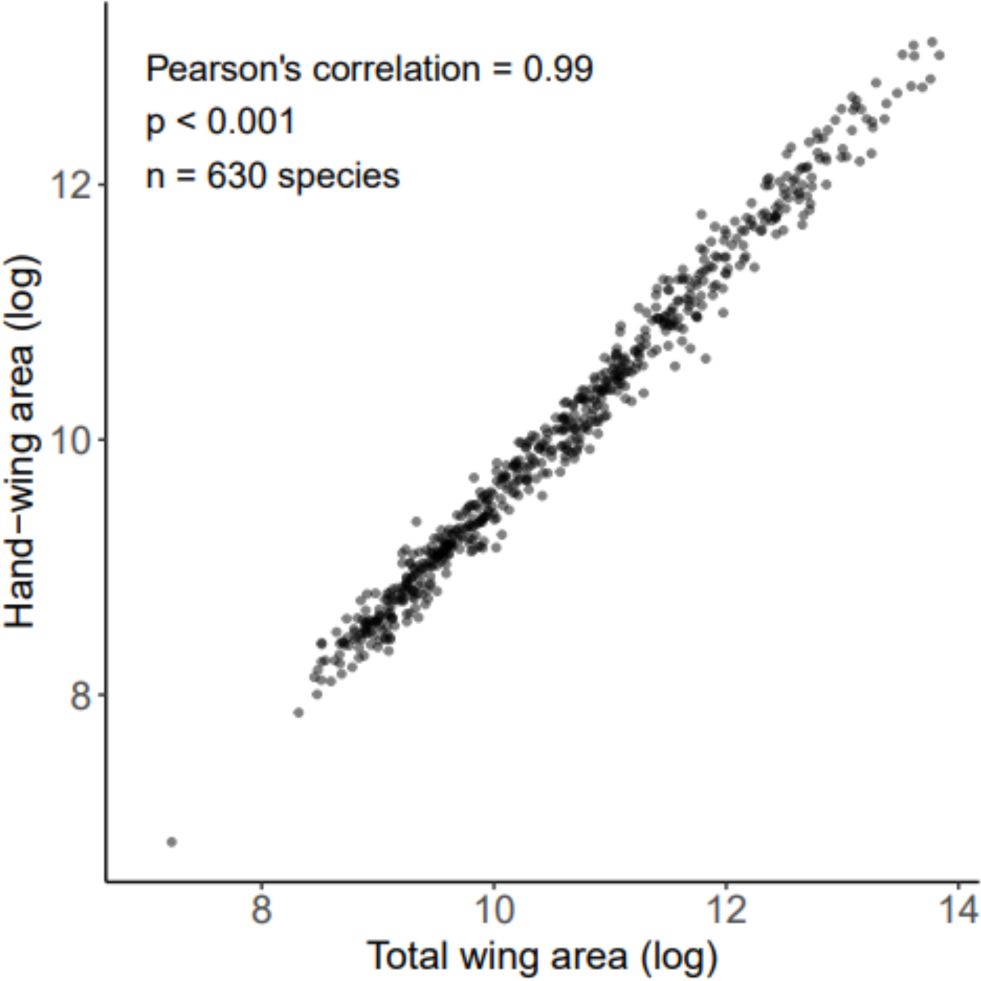
Hand-wing area (HWA) is strongly correlated with the total aeronautical wing area. Total wing area = the aeronautical wing area as defined by most aerodynamic models for aircraft and flying animals, referring to the total area of the underside of both extended wings as well as the underside of the intervening craft or body in between the two wings ^S4^ (Figure S1). We compiled data on total wing area for 630 bird species from published literature estimating total wing area using open-winged museum specimens or live individuals. Full dataset and sources are presented in Data S1. We define HWA as the area of both hand-wings, calculated from wing measurements for the same sample of species ^S5^ using formula WL x SL x π/2 (Figure S1). Both variables (measured in mm^2^) are log-transformed for use in phylogenetic (PGLS) models.

**Figure S3.**
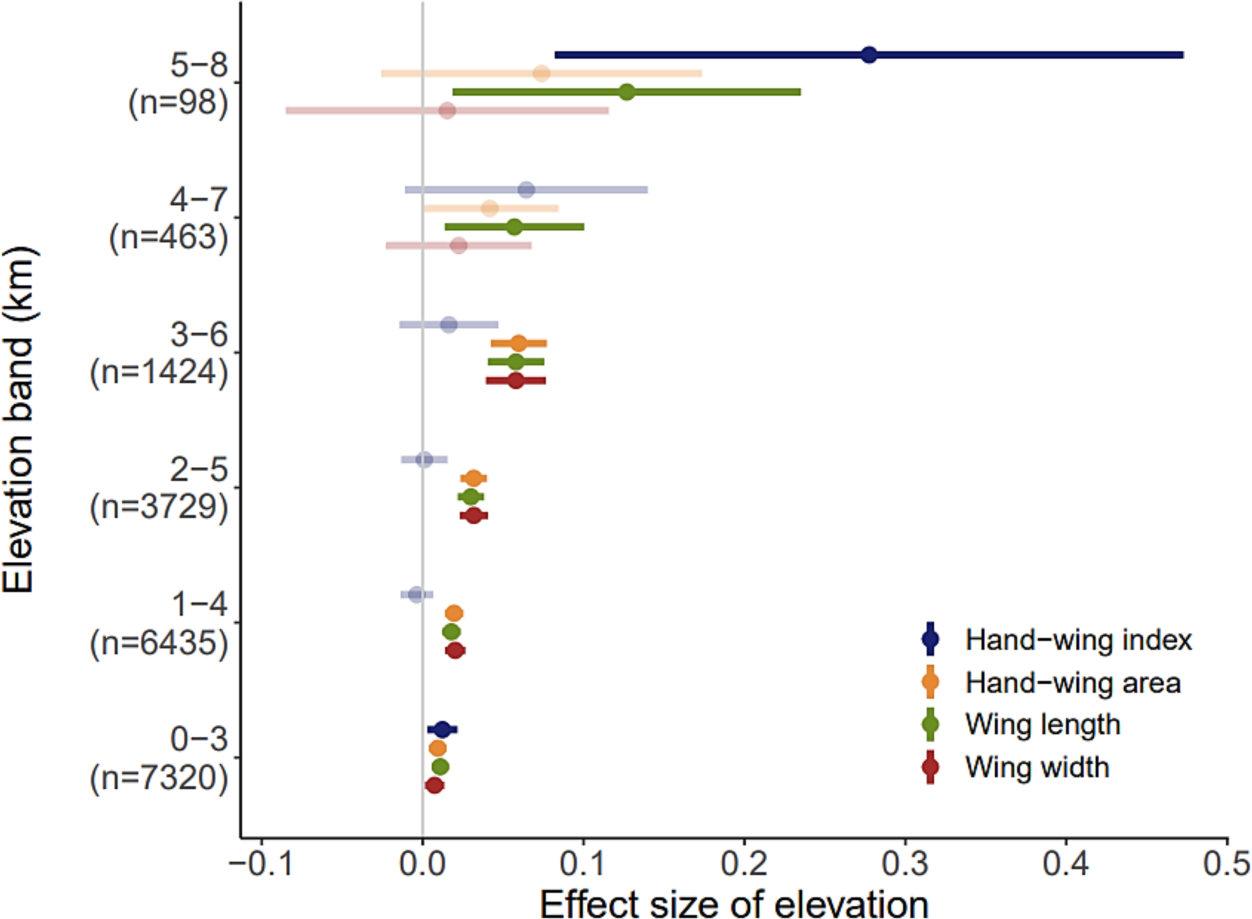
Results of ‘sliding window’ models using non-migratory landbirds. Forest plot shows the parameter estimates of maximum elevation on wing morphology, based on phylogenetic models that sampled species within 3-km wide elevational bands while removing marine and migratory species. The coverage of elevational bands and the corresponding sample sizes are shown on the y-axis. Central points show the mean; error bars show 95% CI. Results were generated using 100 randomly selected trees and averaged via Rubin’s rules. Significant effects are inferred when estimated 95% CI do not span zero (highlighted in a darker shade).

**Figure S4.**
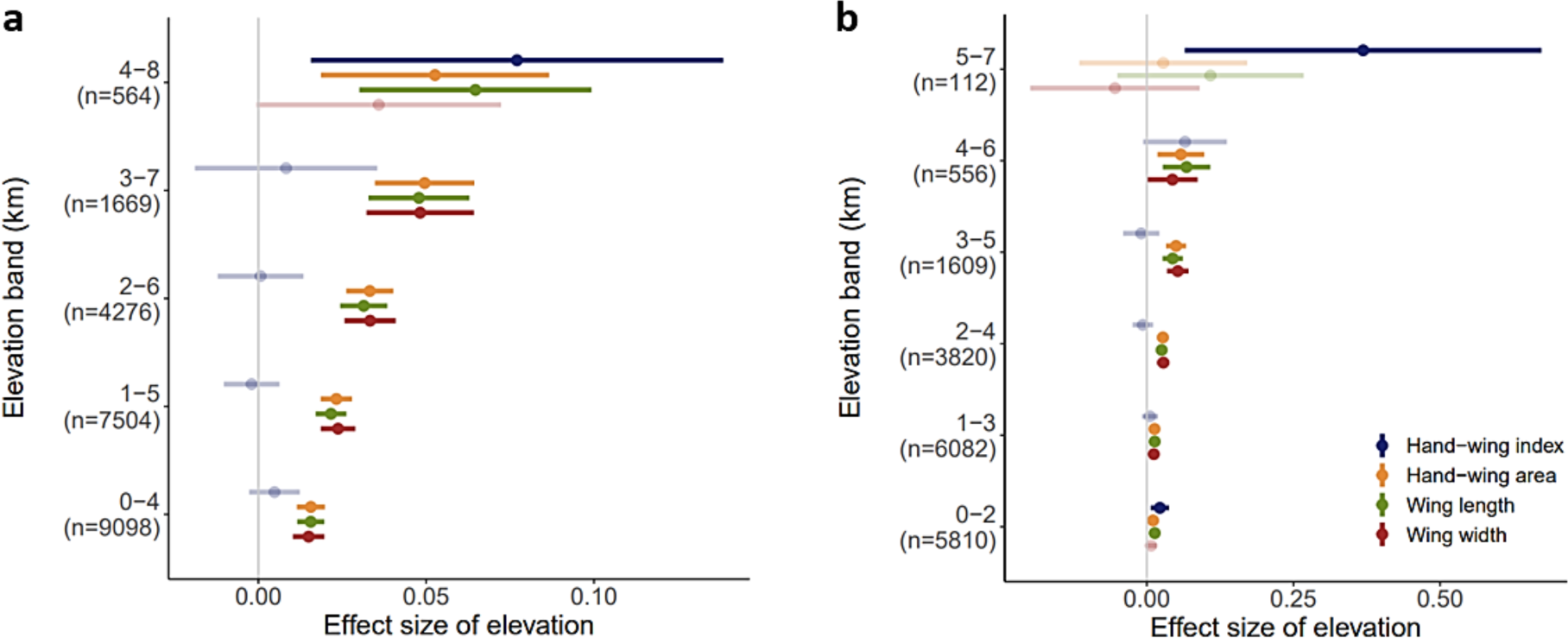
Results of ‘sliding window’ models using alternative bandwidth settings. Forest plot shows the parameter estimates of maximum elevation on wing morphology, based on phylogenetic models that sampled species within 4-km wide (a) or 2-km wide (b) elevational bands. The coverage of elevational bands and the corresponding sample sizes are shown on the y-axis. Central points show the mean; error bars show 95% CI. Results were generated using 100 randomly selected trees and averaged via Rubin’s rules. Significant effects are inferred when estimated 95% CI do not span zero (highlighted in a darker shade). 2-km band models (b) are truncated at 5-7 km because 6-8 km contains only 18 species with limited trait variation which does not support multivariant PGLS models.

**Figure S5.**
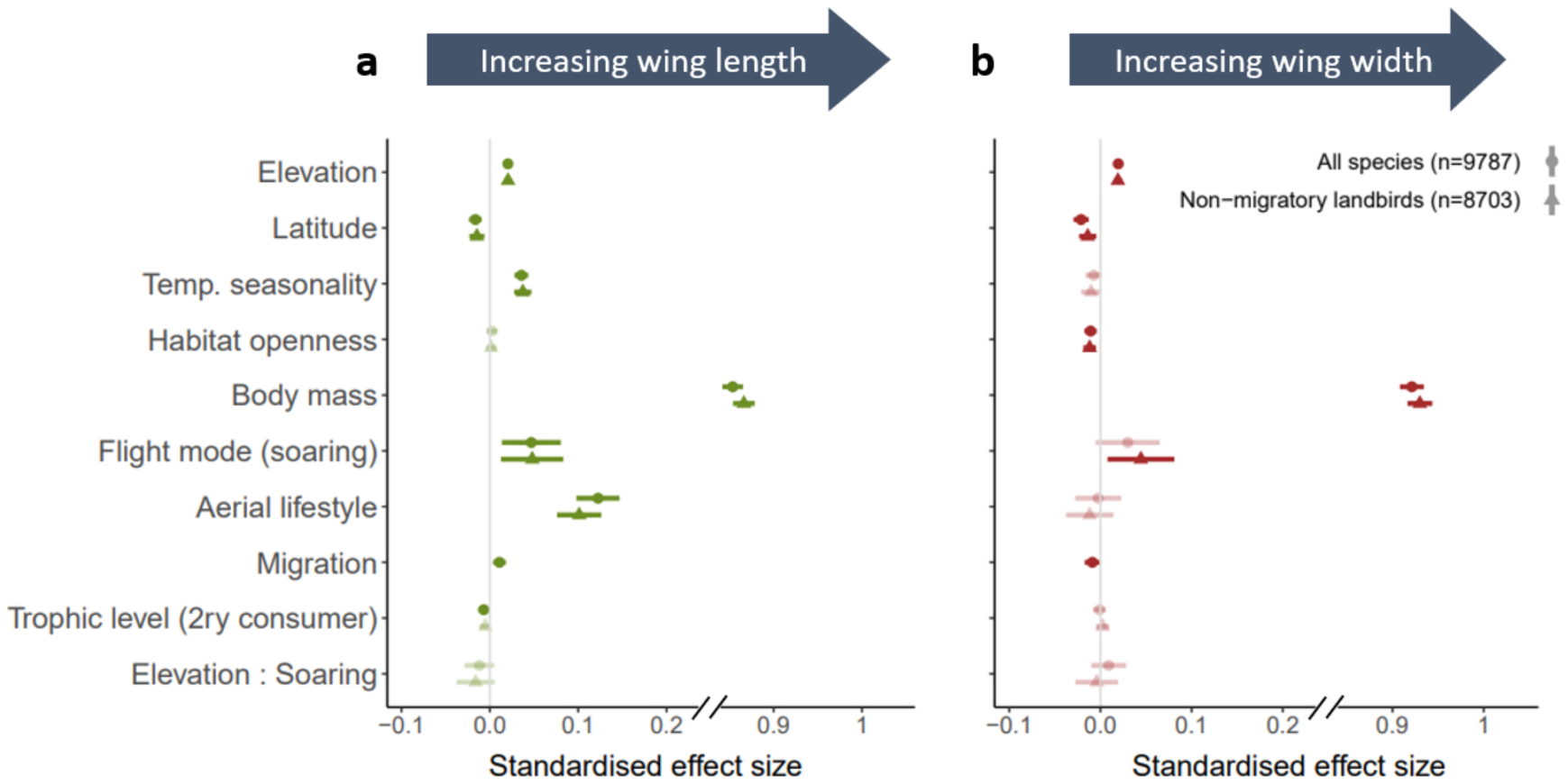
Results of phylogenetic models showing that maximum elevation predicts linear measurements of wing length and wing width. Panels show forest plots with parameter estimates of phylogenetic models testing effects of environmental and ecological factors on (a) wing length and (b) secondary length (see Figure S1). In each case, analyses sampled across all volant birds (n = 9787 species; dots) and non-migratory landbirds only (n = 8703 species; triangles). Central points show the mean; error bars show 95% CI. Results were generated using 100 randomly selected trees and averaged via Rubin’s rules. Positive values indicate a positive correlation between the predictor and the corresponding wing metric. Seabirds and migratory species may be under different selection pressures because of wind-assisted flight and high-elevation migration, respectively, so we repeated models with both these categories of species removed (i.e. restricting to non-migratory landbirds). Significant effects are inferred when estimated 95% CI do not span zero (highlighted in a darker shade).

**Figure S6.**
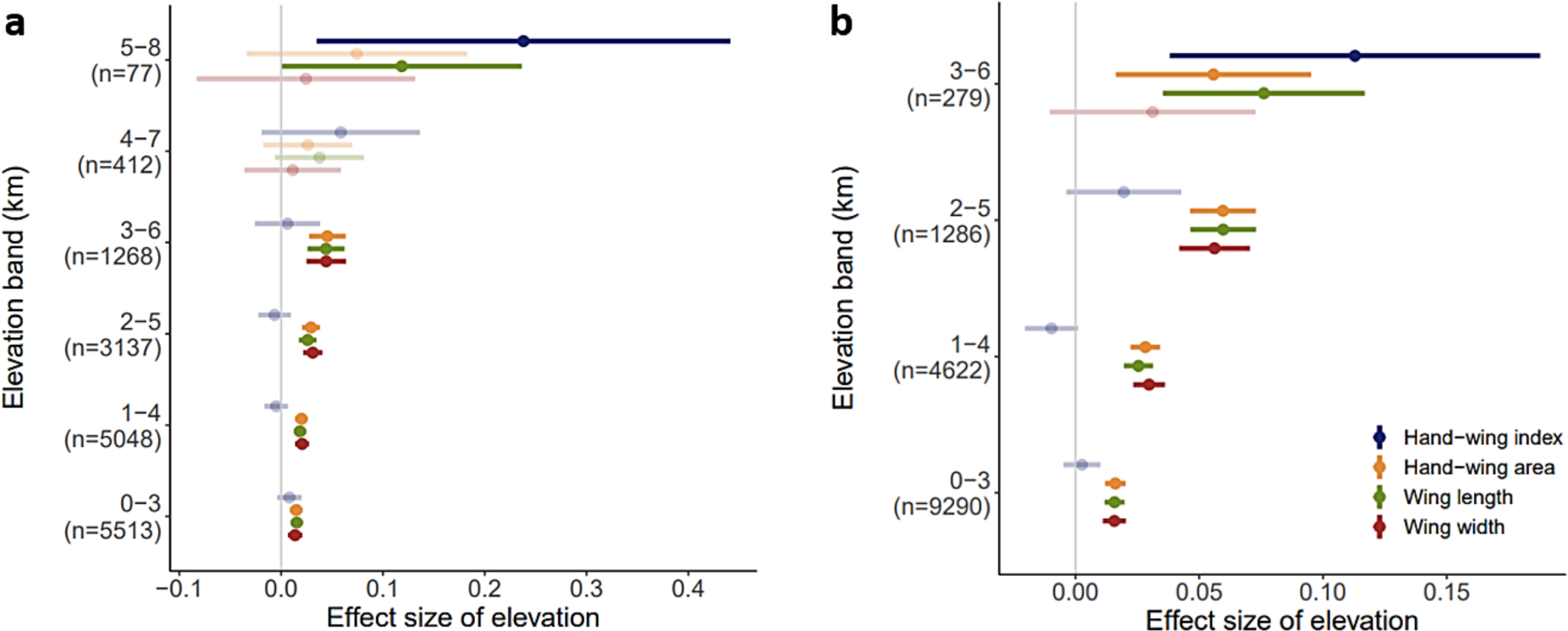
Results of ‘sliding window’ models using species with highest-certainty aerial lifestyle data, with mean elevation as the response variable. Forest plot shows the parameter estimates of elevation on wing morphology based on phylogenetic models that sampled species within 3-km wide elevational bands, using species with highest- certainty aerial lifestyle data (a), or using mean elevation as the response variable (b). The coverage of elevational bands and the corresponding sample sizes are shown on the y-axis. Central points show the mean; error bars show 95% CI. Results were generated using 100 randomly selected trees and averaged via Rubin’s rules. Significant effects are inferred when estimated 95% CI do not span zero (highlighted in a darker shade). In the mean-elevation models (b), species were divided into elevational bands according to their mean elevation instead of maximum elevation in congruence with the model structure (i.e. using mean elevation as the response variable). The highest elevational band in mean-elevation models (b) is 3-6 km because no birds have a mean elevation higher than 6 km in our dataset.

**Figure S7.**
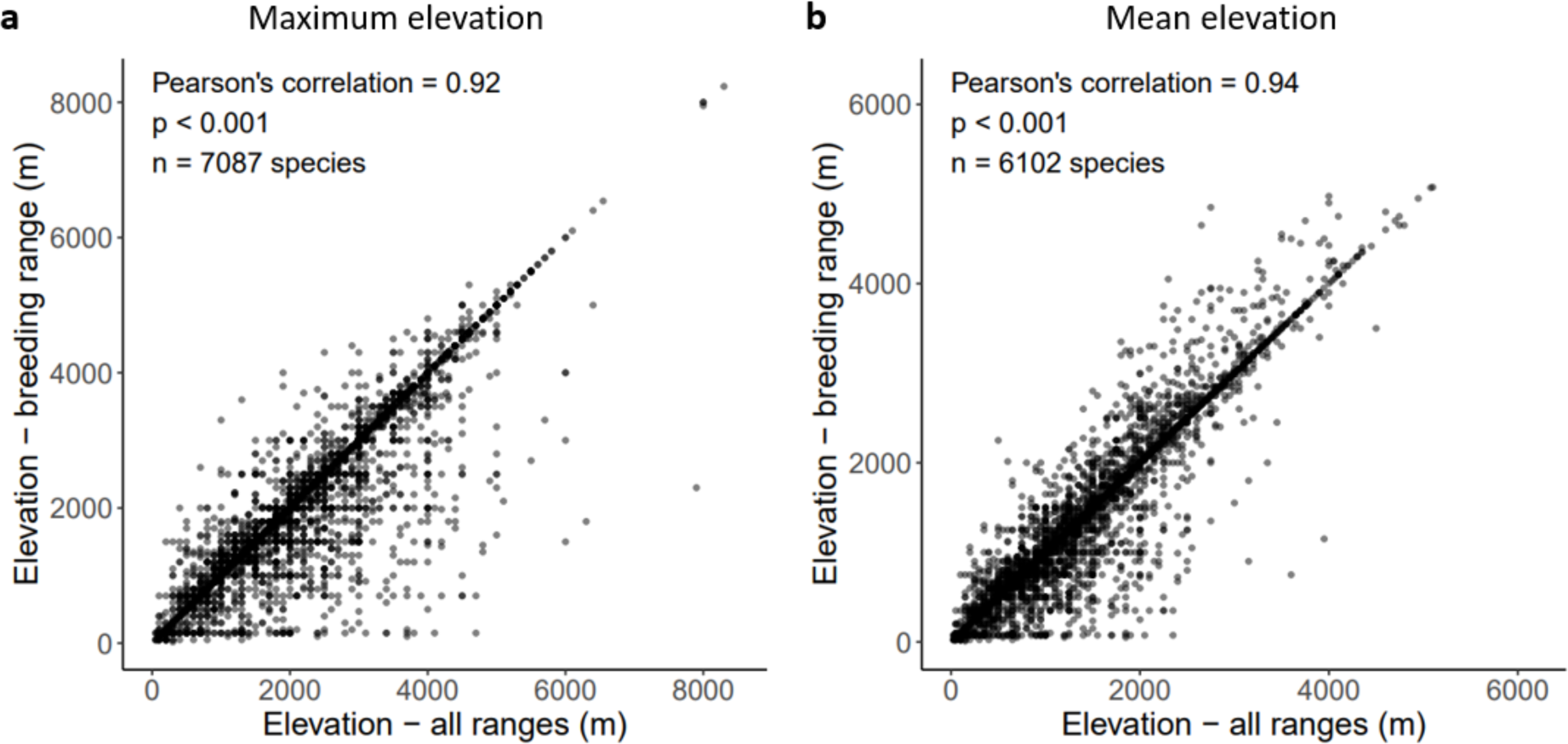
Comparison between elevation data reported by sources using different species ranges. For both maximum (a) and mean (b) elevation, scatterplots show correlations between the elevational data compiled from all known ranges of a species (x axes) and data compiled only from the breeding range (y axes). Data for all-ranges elevation were extracted from ref. ^S6^ and ref. ^S7^; data for breeding-range elevation were extracted from ref. ^S8^. Sampling is restricted to a subset of species for which both types of data are available, including resident species (n = 74% and 70% of sampled species for maximum and mean elevation, respectively) and partial or full migrants (the remaining species).

**Figure S8.**
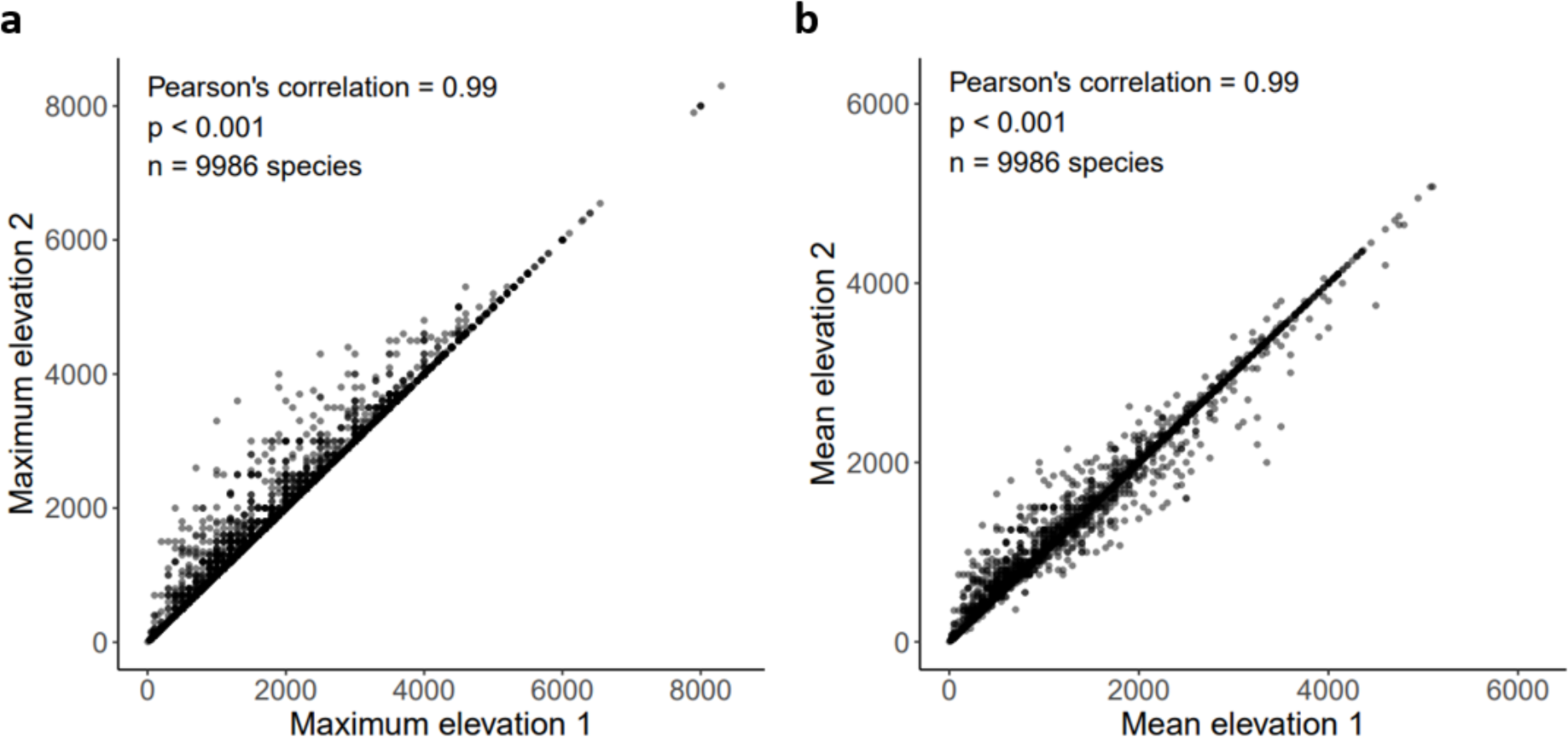
Correlation between different methods for generating elevation data. We collected elevational range data from multiple sources and selected the final elevation data using two alternative methods. Since one major source ^S^^8^ provides elevational range data exclusively from breeding range, increasing uncertainty around the elevational data, we prioritised elevation data that were extracted from ranges throughout the year (including wintering and migration periods) and were published more recently. Thus, when multiple elevation data were available, Method 1 (i.e., Maximum elevation 1, Mean elevation 1) prioritises sources in the order ref. ^S^^6^ > ref. ^S^^7^ > ref. ^S^^8^, such that data from the highest-priority source is used even when it is not the highest elevation. As an alternative measure of elevation, Method 2 (i.e., Maximum elevation 2, Mean elevation 2) uses the highest or lowest elevation reported for the species regardless of the source used (see Methods).

## Supplemental tables

**Table S1.** Effects of predictors on hand-wing index (HWI) estimated from global phylogenetic models. Related to Figure 3. Predictor estimates were generated using 100 randomly selected trees and averaged via Rubin’s rules. For each predictor and summary statistic, we show results using all volant species (top row; n = 9787 species) and non-migratory landbirds only (bottom row; n = 8703 species). CI = confidence interval. The predictor ‘Elevation : Soaring’ shows the estimated effect of the interaction term between elevation and flight mode. The factor ‘Elevation (soaring)’ shows the estimated effect of elevation for soaring species, calculated from the main effect of elevation (Elevation) and the interaction term (Elevation : Soaring). Summary statistics (conditional R^2^ and Pagel’s lambda) were estimated as the mean of 100 conditional R^2^ and Pagel’s lambda values obtained from 100 models. Significant results (those with 95% CI not spanning zero) are highlighted in grey.

| Predictor | Estimate | Std. error | 95% CI (lower) | 95% CI (upper) | $P_t$ value |
| --- | --- | --- | --- | --- | --- |
| (Intercept) | 0.102 | 0.109 | -0.111 | 0.314 | 0.35 |
|  | 0.084 | 0.108 | -0.128 | 0.297 | 0.44 |
| Elevation | 0.007 | 0.003 | 0.000 | 0.013 | < 0.05 |
|  | 0.008 | 0.003 | 0.002 | 0.015 | < 0.05 |
| Latitude | 0.013 | 0.007 | -0.000 | 0.027 | 0.06 |
|  | -0.001 | 0.008 | -0.016 | 0.014 | 0.93 |
| Temp. seasonality | 0.107 | 0.008 | 0.092 | 0.121 | < 0.001 |
|  | 0.119 | 0.009 | 0.102 | 0.137 | < 0.001 |
| Habitat openness | 0.029 | 0.005 | 0.019 | 0.040 | < 0.001 |
|  | 0.029 | 0.006 | 0.018 | 0.040 | < 0.001 |
| Body mass | -0.006 | 0.011 | -0.028 | 0.016 | 0.59 |
|  | -0.000 | 0.012 | -0.023 | 0.022 | 0.98 |
| Flight mode (soaring) | 0.057 | 0.031 | -0.004 | 0.118 | 0.07 |
|  | 0.040 | 0.032 | -0.023 | 0.103 | 0.22 |
| Aerial lifestyle | 0.271 | 0.022 | 0.228 | 0.314 | < 0.001 |
|  | 0.244 | 0.023 | 0.198 | 0.289 | < 0.001 |
| Migration | 0.055 | 0.007 | 0.042 | 0.069 | < 0.001 |
|  | NA | NA | NA | NA | NA |
| Trophic level (2ry consumer) | -0.016 | 0.006 | -0.027 | -0.004 | < 0.01 |
|  | -0.018 | 0.006 | -0.030 | -0.007 | < 0.01 |
| Elevation : Soaring | -0.041 | 0.016 | -0.073 | -0.010 | < 0.01 |
|  | -0.032 | 0.019 | -0.069 | 0.006 | 0.10 |
| Elevation (soaring) * | -0.035 | 0.016 | -0.065 | -0.004 | < 0.05 |
|  | -0.023 | 0.019 | -0.060 | 0.014 | 0.22 |
| **Summary statistics** |  |  |  |  |  |
| Conditional $R^2$ | 0.925 | | | | |
|  | 0.920 |  |  |  |  |
| Pagel's lambda | 0.910 |  |  |  |  |
|  | 0.924 |  |  |  |  |

**Table S2.** Effects of predictors on hand-wing area (HWA) estimated from global phylogenetic models. Related to Figure 3. Predictor estimates were generated using 100 randomly selected trees and averaged via Rubin’s rules. For each predictor and summary statistic, we show results using all volant species (top row; n = 9787 species) and non-migratory landbirds only (bottom row; n = 8703 species). CI = confidence interval. The predictor ‘Elevation : Soaring’ shows the estimated effect of the interaction term between elevation and flight mode. The factor ‘Elevation (soaring)’ shows the estimated effect of elevation for soaring species, calculated from the main effect of elevation (Elevation) and the interaction term (Elevation : Soaring). Summary statistics (conditional R^2^ and Pagel’s lambda) were estimated as the mean of 100 conditional R^2^ and Pagel’s lambda values obtained from 100 models. Significant results (those with 95% CI not spanning zero) are highlighted in grey.

| Predictor | Estimate | Std. error | Lower 95% CI | Higher 95% CI | P<sub>t</sub> value |
| --- | --- | --- | --- | --- | --- |
| (Intercept) | -0.035 | 0.056 | -0.146 | 0.075 | 0.53 |
|  | -0.039 | 0.056 | -0.148 | 0.070 | 0.49 |
| Elevation | 0.021 | 0.002 | 0.017 | 0.024 | < 0.001 |
|  | 0.020 | 0.002 | 0.017 | 0.024 | < 0.001 |
| Latitude | -0.019 | 0.004 | -0.027 | -0.012 | < 0.001 |
|  | -0.015 | 0.004 | -0.023 | -0.006 | < 0.001 |
| Temp. seasonality | 0.015 | 0.004 | 0.007 | 0.023 | < 0.001 |
|  | 0.015 | 0.005 | 0.005 | 0.025 | < 0.01 |
| Habitat openness | -0.004 | 0.003 | -0.010 | 0.002 | 0.22 |
|  | -0.005 | 0.003 | -0.011 | 0.002 | 0.14 |
| Body mass | 0.909 | 0.006 | 0.897 | 0.921 | < 0.001 |
|  | 0.921 | 0.006 | 0.908 | 0.933 | < 0.001 |
| Flight mode (soaring) | 0.039 | 0.017 | 0.007 | 0.072 | < 0.05 |
|  | 0.047 | 0.017 | 0.013 | 0.081 | < 0.01 |
| Aerial lifestyle | 0.066 | 0.012 | 0.043 | 0.090 | < 0.001 |
|  | 0.050 | 0.012 | 0.025 | 0.074 | < 0.001 |
| Migration | 0.002 | 0.004 | -0.006 | 0.009 | 0.65 |
|  | NA | NA | NA | NA | NA |
| Trophic level (2ry consumer) | -0.004 | 0.003 | -0.010 | 0.002 | 0.19 |
|  | -0.002 | 0.003 | -0.008 | 0.005 | 0.65 |
| Elevation : Soaring | -0.002 | 0.009 | -0.019 | 0.015 | 0.81 |
|  | -0.011 | 0.011 | -0.033 | 0.011 | 0.32 |
| Elevation (soaring) * | 0.018 | 0.009 | 0.002 | 0.035 | < 0.05 |
|  | 0.009 | 0.011 | -0.012 | 0.031 | 0.39 |
| **Summary statistics** |  |  |  |  |  |
| Conditional R<sup>2</sup> | 0.977 |  |  |  |  |
|  | 0.976 |  |  |  |  |
| Pagel's lambda | 0.884 |  |  |  |  |
|  | 0.878 |  |  |  |  |

**Table S3.** Effects of predictors on the absolute change in HWI (without body size correction) estimated from global phylogenetic models. Predictor estimates were generated using 100 randomly selected trees and averaged via Rubin’s rules. For each predictor and summary statistic, we show results using all volant species (top row; n = 9787 species) and non-migratory landbirds only (bottom row; n = 8703 species). CI = confidence interval. The predictor ‘Elevation : Soaring’ shows the estimated effect of the interaction term between elevation and flight mode. The factor ‘Elevation (soaring)’ shows the estimated effect of elevation for soaring species, calculated from the main effect of elevation (Elevation) and the interaction term (Elevation : Soaring). Summary statistics (conditional R^2^ and Pagel’s lambda) were estimated as the mean of 100 conditional R^2^ and Pagel’s lambda values obtained from 100 models. Significant results (those with 95% CI not spanning zero) are highlighted in grey.

| Predictor | Estimate | Std. error | Lower 95% CI | Higher 95% CI | P<sub>t</sub> value |
| --- | --- | --- | --- | --- | --- |
| (Intercept) | 0.098 | 0.108 | -0.115 | 0.310 | 0.37 |
|  | 0.084 | 0.108 | -0.127 | 0.296 | 0.44 |
| Elevation | 0.007 | 0.003 | 0.000 | 0.013 | < 0.05 |
|  | 0.008 | 0.003 | 0.002 | 0.015 | < 0.05 |
| Latitude | 0.013 | 0.007 | -0.001 | 0.026 | 0.06 |
|  | -0.001 | 0.008 | -0.016 | 0.014 | 0.93 |
| Temp. seasonality | 0.107 | 0.008 | 0.092 | 0.122 | < 0.001 |
|  | 0.119 | 0.009 | 0.102 | 0.137 | < 0.001 |
| Habitat openness | 0.029 | 0.005 | 0.018 | 0.040 | < 0.001 |
|  | 0.029 | 0.006 | 0.018 | 0.040 | < 0.001 |
| Flight mode (soaring) | 0.056 | 0.031 | -0.005 | 0.117 | 0.07 |
|  | 0.040 | 0.032 | -0.023 | 0.103 | 0.22 |
| Aerial lifestyle | 0.271 | 0.022 | 0.228 | 0.314 | < 0.001 |
|  | 0.244 | 0.023 | 0.198 | 0.289 | < 0.001 |
| Migration | 0.055 | 0.007 | 0.042 | 0.069 | < 0.001 |
|  | NA | NA | NA | NA | NA |
| Trophic level (2ry consumer) | -0.015 | 0.006 | -0.027 | -0.004 | < 0.01 |
|  | -0.018 | 0.006 | -0.030 | -0.007 | < 0.01 |
| Elevation : Soaring | -0.042 | 0.016 | -0.073 | -0.011 | < 0.01 |
|  | -0.032 | 0.019 | -0.069 | 0.006 | 0.10 |
| Elevation (soaring) * | -0.035 | 0.016 | -0.065 | -0.004 | < 0.05 |
|  | -0.023 | 0.019 | -0.060 | 0.014 | 0.22 |
| **Summary statistics** |  |  |  |  |  |
| Conditional R<sup>2</sup> | 0.925 |  |  |  |  |
|  | 0.920 |  |  |  |  |
| Pagel's lambda | 0.910 |  |  |  |  |
|  | 0.924 |  |  |  |  |

**Table S4.** Effects of predictors on the absolute change in HWA (without body size correction) estimated from global phylogenetic models. Predictor estimates were generated using 100 randomly selected trees and averaged via Rubin’s rules. For each predictor and summary statistic, we show results using all volant species (top row; n = 9787 species) and non-migratory landbirds only (bottom row; n = 8703 species). CI = confidence interval. The predictor ‘Elevation : Soaring’ shows the estimated effect of the interaction term between elevation and flight mode. The factor ‘Elevation (soaring)’ shows the estimated effect of elevation for soaring species, calculated from the main effect of elevation (Elevation) and the interaction term (Elevation : Soaring). Summary statistics (conditional R^2^ and Pagel’s lambda) were estimated as the mean of 100 conditional R^2^ and Pagel’s lambda values obtained from 100 models. Significant results (those with 95% CI not spanning zero) are highlighted in grey.

| Predictor | Estimate | Std. error | Lower 95% CI | Higher 95% CI | P<sub>t</sub> value |
| --- | --- | --- | --- | --- | --- |
| (Intercept) | 0.566 | 0.142 | 0.287 | 0.844 | < 0.001 |
|  | 0.567 | 0.143 | 0.286 | 0.848 | < 0.001 |
| Elevation | 0.030 | 0.004 | 0.023 | 0.036 | < 0.001 |
|  | 0.030 | 0.004 | 0.023 | 0.038 | < 0.001 |
| Latitude | 0.021 | 0.007 | 0.007 | 0.034 | < 0.01 |
|  | 0.019 | 0.008 | 0.003 | 0.036 | < 0.05 |
| Temp. seasonality | -0.015 | 0.007 | -0.029 | 0.000 | 0.05 |
|  | -0.007 | 0.010 | -0.026 | 0.011 | 0.44 |
| Habitat openness | 0.020 | 0.006 | 0.009 | 0.031 | < 0.001 |
|  | 0.017 | 0.006 | 0.005 | 0.029 | < 0.01 |
| Flight mode (soaring) | 0.139 | 0.041 | 0.058 | 0.220 | < 0.001 |
|  | 0.141 | 0.046 | 0.051 | 0.230 | < 0.01 |
| Aerial lifestyle | 0.064 | 0.026 | 0.013 | 0.114 | < 0.05 |
|  | 0.050 | 0.028 | -0.005 | 0.104 | 0.07 |
| Migration | -0.006 | 0.007 | -0.020 | 0.007 | 0.37 |
|  | NA | NA | NA | NA | NA |
| Trophic level (2ry consumer) | -0.020 | 0.006 | -0.032 | -0.008 | < 0.01 |
|  | -0.019 | 0.007 | -0.032 | -0.006 | < 0.01 |
| Elevation : Soaring | 0.024 | 0.016 | -0.008 | 0.057 | 0.14 |
|  | 0.054 | 0.022 | 0.012 | 0.096 | < 0.05 |
| Elevation (soaring) * | 0.054 | 0.016 | 0.022 | 0.085 | < 0.001 |
|  | 0.085 | 0.021 | 0.043 | 0.126 | < 0.001 |
| **Summary statistics** |  |  |  |  |  |
| Conditional R<sup>2</sup> | 0.918 |  |  |  |  |
|  | 0.914 |  |  |  |  |
| Pagel's lambda | 0.963 |  |  |  |  |
|  | 0.964 |  |  |  |  |

**Table S5.**
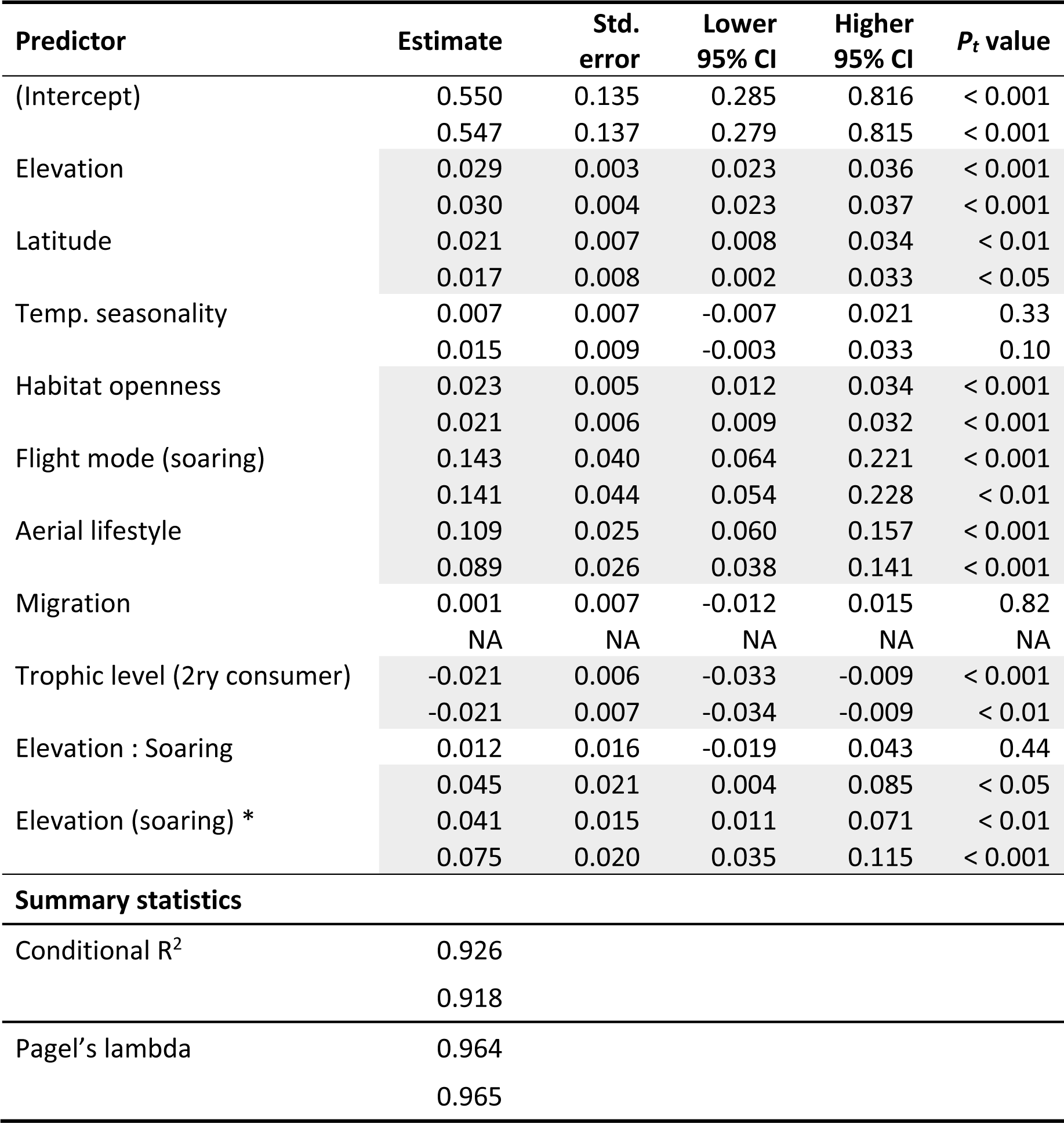
Effects of predictors on the absolute change in wing length (without body size correction) estimated from global phylogenetic models. Predictor estimates were generated using 100 randomly selected trees and averaged via Rubin’s rules. For each predictor and summary statistic, we show results using all volant species (top row; n = 9787 species) and non-migratory landbirds only (bottom row; n = 8703 species). CI = confidence interval. The predictor ‘Elevation : Soaring’ shows the estimated effect of the interaction term between elevation and flight mode. The factor ‘Elevation (soaring)’ shows the estimated effect of elevation for soaring species, calculated from the main effect of elevation (Elevation) and the interaction term (Elevation : Soaring). Summary statistics (conditional R^2^ and Pagel’s lambda) were estimated as the mean of 100 conditional R^2^ and Pagel’s lambda values obtained from 100 models. Significant results (those with 95% CI not spanning zero) are highlighted in grey.

**Table S6.** Effects of predictors on the absolute change in wing width (without body size correction) estimated from global phylogenetic models. Predictor estimates were generated using 100 randomly selected trees and averaged via Rubin’s rules. For each predictor and summary statistic, we show results using all volant species (top row; n = 9787 species) and non-migratory landbirds only (bottom row; n = 8703 species). CI = confidence interval. The predictor ‘Elevation : Soaring’ shows the estimated effect of the interaction term between elevation and flight mode. The factor ‘Elevation (soaring)’ shows the estimated effect of elevation for soaring species, calculated from the main effect of elevation (Elevation) and the interaction term (Elevation : Soaring). Summary statistics (conditional R^2^ and Pagel’s lambda) were estimated as the mean of 100 conditional R^2^ and Pagel’s lambda values obtained from 100 models. Significant results (those with 95% CI not spanning zero) are highlighted in grey.

| Predictor | Estimate | Std. error | Lower 95% CI | Higher 95% CI | P<sub>t</sub> value |
| --- | --- | --- | --- | --- | --- |
| (Intercept) | 0.554 | 0.146 | 0.268 | 0.839 | < 0.001 |
|  | 0.560 | 0.147 | 0.272 | 0.847 | < 0.001 |
| Elevation | 0.029 | 0.004 | 0.021 | 0.036 | < 0.001 |
|  | 0.029 | 0.004 | 0.021 | 0.037 | < 0.001 |
| Latitude | 0.019 | 0.007 | 0.005 | 0.034 | < 0.01 |
|  | 0.020 | 0.009 | 0.003 | 0.037 | < 0.05 |
| Temp. seasonality | -0.037 | 0.008 | -0.052 | -0.021 | < 0.001 |
|  | -0.031 | 0.010 | -0.050 | -0.011 | < 0.01 |
| Habitat openness | 0.015 | 0.006 | 0.004 | 0.027 | < 0.05 |
|  | 0.012 | 0.006 | 0.000 | 0.025 | < 0.05 |
| Flight mode (soaring) | 0.131 | 0.041 | 0.050 | 0.212 | < 0.01 |
|  | 0.136 | 0.046 | 0.046 | 0.225 | < 0.01 |
| Aerial lifestyle | 0.011 | 0.027 | -0.041 | 0.064 | 0.67 |
|  | 0.004 | 0.029 | -0.052 | 0.06 | 0.89 |
| Migration | -0.015 | 0.007 | -0.029 | -0.000 | < 0.05 |
|  | NA | NA | NA | NA | NA |
| Trophic level (2ry consumer) | -0.017 | 0.006 | -0.030 | -0.005 | < 0.01 |
|  | -0.016 | 0.007 | -0.030 | -0.003 | < 0.05 |
| Elevation : Soaring | 0.036 | 0.017 | 0.002 | 0.070 | < 0.05 |
|  | 0.062 | 0.022 | 0.018 | 0.105 | < 0.01 |
| Elevation (soaring) * | 0.065 | 0.017 | 0.032 | 0.098 | < 0.001 |
|  | 0.091 | 0.022 | 0.048 | 0.134 | < 0.001 |
| **Summary statistics** |  |  |  |  |  |
| Conditional R<sup>2</sup> | 0.910 |  |  |  |  |
|  | 0.911 |  |  |  |  |
| Pagel's lambda | 0.959 |  |  |  |  |
|  | 0.962 |  |  |  |  |

**Table S7.** Effects of predictors on HWI estimated from global phylogenetic models, using species with the highest-certainty aerial lifestyle data (n = 6864 species). Predictor estimates were generated using 100 randomly selected trees and averaged via Rubin’s rules. CI = confidence interval. The predictor ‘Elevation : Soaring’ shows the estimated effect of the interaction term between elevation and flight mode. Summary statistics (conditional R^2^ and Pagel’s lambda) were estimated as the mean of 100 conditional R^2^ and Pagel’s lambda values obtained from 100 models. Significant results (those with 95% CI not spanning zero) are highlighted in grey.

| Predictor | Estimate | Std. error | Lower 95% CI | Higher 95% CI | *P*<sub>t</sub> value |
| --- | --- | --- | --- | --- | --- |
| (Intercept) | 0.111 | 0.112 | -0.109 | 0.330 | 0.32 |
| Elevation | 0.004 | 0.004 | -0.004 | 0.012 | 0.32 |
| Latitude | 0.015 | 0.008 | -0.001 | 0.031 | 0.07 |
| Temp. seasonality | 0.102 | 0.009 | 0.084 | 0.119 | < 0.001 |
| Habitat openness | 0.035 | 0.007 | 0.021 | 0.049 | < 0.001 |
| Body mass | -0.018 | 0.013 | -0.044 | 0.008 | 0.18 |
| Flight mode (soaring) | 0.099 | 0.037 | 0.026 | 0.172 | < 0.01 |
| Aerial lifestyle | 0.297 | 0.025 | 0.248 | 0.346 | < 0.001 |
| Migration | 0.048 | 0.008 | 0.032 | 0.064 | < 0.001 |
| Trophic level (2ry consumer) | -0.020 | 0.007 | -0.034 | -0.005 | < 0.01 |
| Elevation : Soaring | -0.047 | 0.018 | -0.082 | -0.012 | < 0.01 |
| **Summary statistics** |  |  |  |  |  |
| Conditional R<sup>2</sup> | 0.932 |  |  |  |  |
| Pagel's lambda | 0.911 |  |  |  |  |

**Table S8.** Effects of predictors on HWI estimated from global phylogenetic models, using mean elevation as the response variable (n = 9787 species). Predictor estimates were generated using 100 randomly selected trees and averaged via Rubin’s rules. CI = confidence interval. The predictor ‘Elevation : Soaring’ shows the estimated effect of the interaction term between elevation and flight mode. Summary statistics (conditional R^2^ and Pagel’s lambda) were estimated as the mean of 100 conditional R^2^ and Pagel’s lambda values obtained from 100 models. Significant results (those with 95% CI not spanning zero) are highlighted in grey.

| Predictor | Estimate | Std. error | Lower 95% CI | Higher 95% CI | P<sub>t</sub> value |
| --- | --- | --- | --- | --- | --- |
| (Intercept) | 0.101 | 0.109 | -0.112 | 0.314 | 0.35 |
| Elevation | 0.002 | 0.003 | -0.005 | 0.008 | 0.61 |
| Latitude | 0.012 | 0.007 | -0.002 | 0.025 | 0.09 |
| Temp. seasonality | 0.109 | 0.007 | 0.094 | 0.124 | < 0.001 |
| Habitat openness | 0.030 | 0.005 | 0.019 | 0.040 | < 0.001 |
| Body mass | -0.005 | 0.011 | -0.027 | 0.016 | 0.62 |
| Flight mode (soaring) | 0.054 | 0.031 | -0.007 | 0.115 | 0.08 |
| Aerial lifestyle | 0.271 | 0.022 | 0.228 | 0.314 | < 0.001 |
| Migration | 0.055 | 0.007 | 0.041 | 0.068 | < 0.001 |
| Trophic level (2ry consumer) | -0.016 | 0.006 | -0.027 | -0.004 | < 0.01 |
| Elevation : Soaring | -0.044 | 0.020 | -0.083 | -0.005 | < 0.05 |
| **Summary statistics** |  |  |  |  |  |
| Conditional R<sup>2</sup> | 0.924 |  |  |  |  |
| Pagel's lambda | 0.910 |  |  |  |  |

**Table S9.** Effects of predictors on HWA estimated from global phylogenetic models, using species with the highest-certainty aerial lifestyle data (n = 6864 species). Predictor estimates were generated using 100 randomly selected trees and averaged via Rubin’s rules. CI = confidence interval. The predictor ‘Elevation : Soaring’ shows the estimated effect of the interaction term between elevation and flight mode. Summary statistics (conditional R^2^ and Pagel’s lambda) were estimated as the mean of 100 conditional R^2^ and Pagel’s lambda values obtained from 100 models. Significant results (those with 95% CI not spanning zero) are highlighted in grey.

| Predictor | Estimate | Std. error | Lower 95% CI | Higher 95% CI | *P*<sub>t</sub> value |
| --- | --- | --- | --- | --- | --- |
| (Intercept) | -0.059 | 0.057 | -0.171 | 0.054 | 0.31 |
| Elevation | 0.022 | 0.002 | 0.018 | 0.026 | < 0.001 |
| Latitude | -0.020 | 0.004 | -0.029 | -0.012 | < 0.001 |
| Temp. seasonality | 0.014 | 0.005 | 0.005 | 0.023 | < 0.01 |
| Habitat openness | -0.002 | 0.004 | -0.009 | 0.006 | 0.63 |
| Body mass | 0.925 | 0.007 | 0.911 | 0.939 | < 0.001 |
| Flight mode (soaring) | 0.058 | 0.019 | 0.021 | 0.095 | < 0.01 |
| Aerial lifestyle | 0.075 | 0.013 | 0.049 | 0.100 | < 0.001 |
| Migration | 0.007 | 0.004 | -0.002 | 0.015 | 0.12 |
| Trophic level (2ry consumer) | -0.003 | 0.004 | -0.010 | 0.004 | 0.43 |
| Elevation : Soaring | -0.008 | 0.009 | -0.026 | 0.010 | 0.40 |
| **Summary statistics** |  |  |  |  |  |
| Conditional R<sup>2</sup> | 0.980 |  |  |  |  |
| Pagel's lambda | 0.899 |  |  |  |  |

**Table S10.** Effects of predictors on HWA estimated from global phylogenetic models, using mean elevation as the response variable (n = 9787 species). Predictor estimates were generated using 100 randomly selected trees and averaged via Rubin’s rules. CI = confidence interval. The predictor ‘Elevation : Soaring’ shows the estimated effect of the interaction term between elevation and flight mode. Summary statistics (conditional R^2^ and Pagel’s lambda) were estimated as the mean of 100 conditional R^2^ and Pagel’s lambda values obtained from 100 models. Significant results (those with 95% CI not spanning zero) are highlighted in grey.

| Predictor | Estimate | Std. error | Lower 95% CI | Higher 95% CI | *P*<sub>t</sub> value |
| --- | --- | --- | --- | --- | --- |
| (Intercept) | -0.034 | 0.056 | -0.145 | 0.076 | 0.54 |
| Elevation | 0.022 | 0.002 | 0.019 | 0.026 | < 0.001 |
| Latitude | -0.018 | 0.004 | -0.026 | -0.011 | < 0.001 |
| Temp. seasonality | 0.017 | 0.004 | 0.009 | 0.025 | < 0.001 |
| Habitat openness | -0.004 | 0.003 | -0.010 | 0.002 | 0.22 |
| Body mass | 0.908 | 0.006 | 0.896 | 0.919 | < 0.001 |
| Flight mode (soaring) | 0.040 | 0.017 | 0.007 | 0.072 | < 0.05 |
| Aerial lifestyle | 0.067 | 0.012 | 0.043 | 0.090 | < 0.001 |
| Migration | 0.002 | 0.004 | -0.006 | 0.010 | 0.60 |
| Trophic level (2ry consumer) | -0.004 | 0.003 | -0.010 | 0.002 | 0.19 |
| Elevation : Soaring | 0.002 | 0.011 | -0.020 | 0.023 | 0.87 |
| **Summary statistics** |  |  |  |  |  |
| Conditional R<sup>2</sup> | 0.977 |  |  |  |  |
| Pagel's lambda | 0.884 |  |  |  |  |

## References

1. Weeks, B.C., O’Brien, B.K., Chu, J.J., Claramunt, S., Sheard, C., and Tobias, J.A. (2022). Morphological adaptations linked to flight efficiency and aerial lifestyle determine natal dispersal distance in birds. Funct. Ecol. 36, 1681–1689. 10.1111/1365-2435.14056.

2. Sekar, S. (2012). A meta-analysis of the traits affecting dispersal ability in butterflies: can wingspan be used as a proxy? J. Anim. Ecol. 81, 174–184. 10.1111/J.1365-2656.2011.01909.X.

3. Luo, B., Santana, S.E., Pang, Y., Wang, M., Xiao, Y., and Feng, J. (2019). Wing morphology predicts geographic range size in vespertilionid bats. Sci. Rep. 9, 4526. 10.1038/s41598-019-41125-0.

4. Wootton, R.J. (1991). The functional morphology of the wings of Odonata. Adv. Odonatol. 5, 153–169.

5. Hamilton, T.H. (1961). The adaptive significances of intraspecific trends of variation in wing length and body size among bird species. Evolution 15, 180–195. 10.1111/J.1558-5646.1961.TB03142.X.

6. Marchetti, K., Price, T., and Richman, A. (1995). Correlates of wing morphology with foraging behaviour and migration distance in the genus Phylloscopus. J. Avian. Biol. 26, 177–181. 10.2307/3677316.

7. Norberg, U.M., and Rayner, J.M. V. (1987). Ecological morphology and flight in bats (Mammalia; Chiroptera): wing adaptations, flight performance, foraging strategy and echolocation. Philos. Trans. R. Soc. Lond. B Biol. Sci. 316, 335–427. 10.1098/RSTB.1987.0030.

8. 8. Le Roy, C., Debat, V., and Llaurens, V. (2019). Adaptive evolution of butterfly wing shape: from morphology to behaviour. Biol. Rev. 94, 1261–1281. 10.1111/BRV.12500.

9. Sheard, C., Neate-Clegg, M.H.C., Alioravainen, N., Jones, S.E.I., Vincent, C., MacGregor, H.E.A., Bregman, T.P., Claramunt, S., and Tobias, J.A. (2020). Ecological drivers of global gradients in avian dispersal inferred from wing morphology. Nat. Commun. 11, 2463. 10.1038/s41467-020-16313-6.

10. Outomuro, D., Golab, M.J., Johansson, F., and Sniegula, S. (2021). Body and wing size, but not wing shape, vary along a large-scale latitudinal gradient in a damselfly. Sci. Rep. 11, 18642. 10.1038/s41598-021-97829-9.

11. ICAO (1993). Manual of the ICAO Standard Atmosphere: extended to 80 kilometres (262,500 feet) (International Civil Aviation Organization. Montreal, Canada).

12. Hedenström, A. (2002). Aerodynamics, evolution and ecology of avian flight. Trends Ecol. Evol. 17, 415–422. 10.1016/S0169-5347(02)02568-5.

13. Norberg, U.M. (1990). Vertebrate flight: mechanics, physiology, morphology, ecology and evolution (Springer, Berlin, Heidelberg). 10.1007/978-3-642-83848-4.

14. Youngflesh, C., Saracco, J.F., Siegel, R.B., and Tingley, M.W. (2022). Abiotic conditions shape spatial and temporal morphological variation in North American birds. Nat. Ecol. Evol. 6, 1860–1870. 10.1038/s41559-022-01893-x.

15. Feinsinger, P., Colwell, R.K., Terborgh, J., and Chaplin, S.B. (1979). Elevation and the morphology, flight energetics, and foraging ecology of tropical hummingbirds. Am. Nat. 113, 481–497. 10.1086/283408.

16. Altshuler, D.L., and Dudley, R. (2006). The physiology and biomechanics of avian flight at high altitude. Integr. Comp. Biol. 46, 62–71. 10.1093/ICB/ICJ008.

17. Keller, I., Alexander, J.M., Holderegger, R., and Edwards, P.J. (2013). Widespread phenotypic and genetic divergence along altitudinal gradients in animals. J. Evol. Biol. 26, 2527–2543. 10.1111/JEB.12255.

18. Dillon, M.E., Frazier, M.R., and Dudley, R. (2006). Into thin air: Physiology and evolution of alpine insects. Integr. Comp. Biol. 46, 49–61. 10.1093/ICB/ICJ007.

19. Claramunt, S., Derryberry, E.P., Remsen, J. V., and Brumfield, R.T. (2012). High dispersal ability inhibits speciation in a continental radiation of passerine birds. Proc. Biol. Sci. 279, 1567– 1574. 10.1098/RSPB.2011.1922.

20. Pigot, A.L., Jetz, W., Sheard, C., and Tobias, J.A. (2018). The macroecological dynamics of species coexistence in birds. Nat. Ecol. Evol. 2, 1112–1119. 10.1038/s41559-018-0572-9.

21. Weeks, B.C., Willard, D.E., Zimova, M., Ellis, A.A., Witynski, M.L., Hennen, M., and Winger, B.M. (2020). Shared morphological consequences of global warming in North American migratory birds. Ecol. Lett. 23, 316–325. 10.1111/ELE.13434.

22. Weeks, T.L., Betts, M.G., Pfeifer, M., Wolf, C., Banks-Leite, C., Barbaro, L., Barlow, J., Cerezo, A., Kennedy, C.M., Kormann, U.G., et al. (2023). Climate-driven variation in dispersal ability predicts responses to forest fragmentation in birds. Nat. Ecol. Evol. 7, 1079–1091. 10.1038/s41559-023-02077-x.

23. Gilroy, J.J., Gill, J.A., Butchart, S.H.M., Jones, V.R., and Franco, A.M.A. (2016). Migratory diversity predicts population declines in birds. Ecol. Lett. 19, 308–317. 10.1111/ELE.12569.

24. Jønsson, K.A., Tøttrup, A.P., Borregaard, M.K., Keith, S.A., Rahbek, C., and Thorup, K. (2016). Tracking animal dispersal: From individual movement to community assembly and global range dynamics. Trends Ecol. Evol. 31, 204–214. 10.1016/j.tree.2016.01.003.

25. Straus, S., Forbes, C., Little, C.J., Germain, R.M., Main, D.A., O’Connor, M.I., Thompson, P.L., Ford, A.T., Gravel, D., and Guzman, L.M. (2024). Macroecological constraints on species’ ‘movement profiles’: Body mass does not explain it all. Glob. Ecol. Biogeogr. 33, 227–243. 10.1111/GEB.13786.

26. Winger, B.M., Auteri, G.G., Pegan, T.M., and Weeks, B.C. (2019). A long winter for the Red Queen: rethinking the evolution of seasonal migration. Biol. Rev. 94, 737–752. 10.1111/BRV.12476.

27. 27. Jocque, M., Field, R., Brendonck, L., and De Meester, L. (2010). Climatic control of dispersal– ecological specialization trade-offs: a metacommunity process at the heart of the latitudinal diversity gradient? Glob. Ecol. Biogeogr. 19, 244–252. 10.1111/J.1466-8238.2009.00510.X.

28. Pennycuick, C.J. (2008). Modelling the flying bird (Elsevier, Amsterdam).

29. Mayr, E. (1963). Animal species and evolution (Harvard University Press, Cambridge, Massachusetts).

30. Hull, D.G. (2007). Fundamentals of airplane flight mechanics (Springer, Berlin).

31. Segre, P.S., Dakin, R., Read, T.J.G., Straw, A.D., and Altshuler, D.L. (2016). Mechanical constraints on flight at high elevation decrease maneuvering performance of hummingbirds. Curr. Biol. 26, 3368–3374. 10.1016/J.CUB.2016.10.028.

32. Baldwin, M.W., Winkler, H., Organ, C.L., and Helm, B. (2010). Wing pointedness associated with migratory distance in common-garden and comparative studies of stonechats (*Saxicola torquata*). J. Evol. Biol. 23, 1050–1063. 10.1111/J.1420-9101.2010.01975.X.

33. García, J., Arizaga, J., Rodríguez, J.I., Alonso, D., and Suárez-Seoane, S. (2021). Morphological differentiation in a migratory bird across geographic gradients in mountains of southern Europe. J. Biogeogr. 48, 2828–2838. 10.1111/JBI.14242.

34. Sun, Y.F., Ren, Z.P., Wu, Y.F., Lei, F.M., Dudley, R., and Li, D.M. (2016). Flying high: limits to flight performance by sparrows on the Qinghai-Tibet Plateau. J. Exp. Biol. 219, 3642–3648. 10.1242/jeb.142216.

35. Rand, A.L. (1936). Altitudinal variation in New Guinea birds. Am. Mus. Novit. 890, 1–14. https://digitallibrary.amnh.org/items/84d17d07-7216-4412-8c42-c33dbdd591b3.

36. Traylor, M.A. (1950). Altitudinal variation in Bolivian birds. Condor 52, 123–126. 10.2307/1364896.

37. Moreau, R.E. (1957). Variation in the western Zosteropidae (Aves). Bull. Br. Mus. Nat. Hist. Zool. 4, 309–433. https://www.biodiversitylibrary.org/page/26203580.

38. Stresemann, E. (1941). Bemerkungen über *Zonotricha capensi*. Ornithologische Monatsberichte 49, 60–61.

39. Bears, H., Drever, M.C., Martin, K., Bears, H., Drever, M.C., and Martin, K. (2008). Comparative morphology of dark-eyed juncos *Junco hyemalis* breeding at two elevations: a common aviary experiment. J. Avian. Biol. 39, 152–162. 10.1111/J.2008.0908-8857.04191.X.

40. Gutiérrez-Pinto, N., McCracken, K.G., Alza, L., Tubaro, P., Kopuchian, C., Astie, A., and Cadena, C.D. (2014). The validity of ecogeographical rules is context-dependent: testing for Bergmann’s and Allen’s rules by latitude and elevation in a widespread Andean duck. Biol. J. Linn. Soc. Lond. 111, 850–862. 10.1111/bij.12249.

41. Sander, M.M., and Chamberlain, D. (2020). Evidence for intraspecific phenotypic variation in songbirds along elevation gradients in central Europe. Ibis 162, 1355–1362. 10.1111/IBI.12843.

42. Lee, S.Y., Scott, G.R., and Milsom, W.K. (2008). Have wing morphology or flight kinematics evolved for extreme high altitude migration in the bar-headed goose? Comp. Biochem. Physiol. C Toxicol. Pharmacol. 148, 324–331. 10.1016/J.CBPC.2008.05.009.

43. Boyce, A.J., Shakya, S., Sheldon, F.H., Moyle, R.G., and Martin, T.E. (2019). Biotic interactions are the dominant drivers of phylogenetic and functional structure in bird communities along a tropical elevational gradient. Auk 136, ukz054. 10.1093/AUK/UKZ054.

44. Ceresa, F., Vitulano, S., Pes, M., Tomasi, L., Brambilla, M., Kvist, L., Pedrini, P., Anderle, M., Hilpold, A., and Kranebitter, P. (2022). Variation in wing morphology is related to breeding environment in a high-elevation specialist bird. J. Avian. Biol. 2022, e03007. 10.1111/JAV.03007.

45. Tiede, Y., Hemp, C., Schmidt, A., Nauss, T., Farwig, N., and Brandl, R. (2018). Beyond body size: consistent decrease of traits within orthopteran assemblages with elevation. Ecology 99, 2090–2102. 10.1002/ECY.2436.

46. Rendoll-Cárcamo, J., Gañán, M., Madriz, R.I., Convey, P., and Contador, T. (2023). Wing reduction and body size variation along a steep elevation gradient: a case study with Magellanic sub-Antarctic mayflies and stoneflies. Front. Ecol. Evol. 11, 1188889. 10.3389/fevo.2023.1188889.

47. Hodkinson, I.D. (2005). Terrestrial insects along elevation gradients: species and community responses to altitude. Biol. Rev. 80, 489–513. 10.1017/S1464793105006767.

48. Scacco, M., Flack, A., Duriez, O., Wikelski, M., and Safi, K. (2019). Static landscape features predict uplift locations for soaring birds across Europe. R. Soc. Open. Sci. 6, 181440. 10.1098/RSOS.181440.

49. Tobias, J.A., Sheard, C., Pigot, A.L., Devenish, A.J.M., Yang, J., Sayol, F., Neate-Clegg, M.H.C., Alioravainen, N., Weeks, T.L., Barber, R.A., et al. (2022). AVONET: morphological, ecological and geographical data for all birds. Ecol. Lett. 25, 581–597. 10.1111/ele.13898.

50. Wright, N.A., Gregory, T.R., and Witt, C.C. (2014). Metabolic ‘engines’ of flight drive genome size reduction in birds. Proc. Biol. Sci. *281*, 20132780. 10.1098/RSPB.2013.2780.

51. Stoddard, M.C., Yong, E.H., Akkaynak, D., Sheard, C., Tobias, J.A., and Mahadevan, L. (2017). Avian egg shape: Form, function, and evolution. Science 356, 1249–1254. 10.1126/science.aaj1945.

52. Lockwood, R., Swaddle, J.P., and Rayner, J.M. V. (1998). Avian wingtip shape reconsidered: wingtip shape indices and morphological adaptations to migration. J. Avian. Biol. 29, 273–292. 10.2307/3677110.

53. Fu, H., Su, M., Chu, J.J., Margaritescu, A., and Claramunt, S. (2023). New methods for estimating the total wing area of birds. Ecol. Evol. 13, e10480. 10.1002/ECE3.10480.

54. Hedenström, A. (1993). Migration by soaring or flapping flight in birds: the relative importance of energy cost and speed. Philos. Trans. R. Soc. Lond. B Biol. Sci. 342, 353–361. 10.1098/RSTB.1993.0164.

55. Paradis, E., Baillie, S.R., Sutherland, W.J., and Gregory, R.D. (1998). Patterns of natal and breeding dispersal in birds. J. Anim. Ecol. 67, 518–536. 10.1046/J.1365-2656.1998.00215.X.

56. Jenkins, S.H. (1981). Common patterns in home range-body size relationships of birds and mammals. Am. Nat. 118, 126–128. 10.1086/283807.

57. Sachs, G. (2005). Minimum shear wind strength required for dynamic soaring of albatrosses. Ibis 147, 1–10. 10.1111/J.1474-919X.2004.00295.X.

58. Lindström, Å., Alerstam, T., Andersson, A., Bäckman, J., Bahlenberg, P., Bom, R., Ekblom, R., Klaassen, R.H.G., Korniluk, M., Sjöberg, S., et al. (2021). Extreme altitude changes between night and day during marathon flights of great snipes. Curr. Biol. 31, 3433–3439.e3. 10.1016/J.CUB.2021.05.047.

59. Sjöberg, S., Malmiga, G., Nord, A., Andersson, A., Bäckman, J., Tarka, M., Willemoes, M., Thorup, K., Hansson, B., Alerstam, T., et al. (2021). Extreme altitudes during diurnal flights in a nocturnal songbird migrant. Science 372, 646–648. 10.1126/science.abe7291.

60. Schachner, E.R., Moore, A.J., Martinez, A., Diaz, R.E., Echols, M.S., Atterholt, J., W. P. Kissane, R., Hedrick, B.P., and Bates, K.T. (2024). The respiratory system influences flight mechanics in soaring birds. Nature 630, 671–676. 10.1038/s41586-024-07485-y.

61. Parmesan, C. (2006). Ecological and evolutionary responses to recent climate change. Annu. Rev. Ecol. Evol. Syst. 37, 637–669. 10.1146/ANNUREV.ECOLSYS.37.091305.110100.

62. Neate-Clegg, M.H.C., Jones, S.E.I., Tobias, J.A., Newmark, W.D., and Şekercioǧlu, Ç.H. (2021). Ecological correlates of elevational range shifts in tropical birds. Front. Ecol. Evol. 9, 621749. 10.3389/fevo.2021.621749.

63. Lenoir, J., Bertrand, R., Comte, L., Bourgeaud, L., Hattab, T., Murienne, J., and Grenouillet, G. (2020). Species better track climate warming in the oceans than on land. Nat. Ecol. Evol. 4, 1044–1059. 10.1038/s41559-020-1198-2.

64. Spence, A.R., and Tingley, M.W. (2020). The challenge of novel abiotic conditions for species undergoing climate-induced range shifts. Ecography 43, 1571–1590. 10.1111/ECOG.05170.

65. Colwell, R.K., Brehm, G., Cardelús, C.L., Gilman, A.C., and Longino, J.T. (2008). Global warming, elevational range shifts, and lowland biotic attrition in the wet tropics. Science 322, 258–261. 10.1126/science.1162547.

66. Wright, S.J., Muller-Landau, H.C., and Schipper, J. (2009). The future of tropical species on a warmer planet. Conserv. Biol. 23, 1418–1426. 10.1111/J.1523-1739.2009.01337.X.

67. Jankowski, J.E., Robinson, S.K., and Levey, D.J. (2010). Squeezed at the top: Interspecific aggression may constrain elevational ranges in tropical birds. Ecology 91, 1877–1884. 10.1890/09-2063.1.

68. Freeman, B.G., Tobias, J.A., and Schluter, D. (2019). Behavior influences range limits and patterns of coexistence across an elevational gradient in tropical birds. Ecography 42, 1832– 1840. 10.1111/ECOG.04606.

69. Projecto-Garcia, J., Natarajan, C., Moriyama, H., Weber, R.E., Fago, A., Cheviron, Z.A., Dudley, R., McGuire, J.A., Witt, C.C., and Storz, J.F. (2013). Repeated elevational transitions in hemoglobin function during the evolution of Andean hummingbirds. Proc. Natl. Acad. Sci. 110, 20669–20674. 10.1073/pnas.1315456110.

70. Martin, K., Zwaan, D.R. de, Scridel, D., and Altamirano, T.A. (2023). Avian adaptations to high mountain habitats. In Ecology and Conservation of Mountain Birds (Cambridge University Press, Cambridge), pp. 35–89. 10.1017/9781108938570.003.

71. Ivy, C.M., and Williamson, J.L. (2024). On the physiology of high-altitude flight and altitudinal migration in birds. Integr. Comp. Biol., icae062. 10.1093/ICB/ICAE062.

72. Moore, M.P., Shaich, J., and Stroud, J.T. (2023). Upslope migration is slower in insects that depend on metabolically demanding flight. Nat. Clim. Chang. 13, 1063–1066. 10.1038/s41558-023-01794-2.

73. Tobias, J.A., Sheard, C., Seddon, N., Meade, A., Cotton, A.J., and Nakagawa, S. (2016). Territoriality, social bonds, and the evolution of communal signaling in birds. Front. Ecol. Evol. 4, 74. 10.3389/fevo.2016.00074.

74. Baliga, B., Szabo, I., and Altshuler, D.L. (2019). Range of motion in the avian wing is strongly associated with flight behavior and body mass. Sci. Adv. 5, eaaw6670. 10.1126/sciadv.aaw6670.

75. Sherub, S., Bohrer, G., Wikelski, M., and Weinzierl, R. (2016). Behavioural adaptations to flight into thin air. Biol. Lett. 12, 20160432. 10.1098/RSBL.2016.0432.

76. Harvey, C., Baliga, V.B., Wong, J.C.M., Altshuler, D.L., and Inman, D.J. (2022). Birds can transition between stable and unstable states via wing morphing. Nature 603, 648–653. 10.1038/s41586-022-04477-8.

77. Halperin, D.J., Guinn, T.A., Strazzo, S.E., and Thomas, R.L. (2022). Density altitude: climatology of daily maximum values and evaluation of approximations for general aviation. Weather Clim. Soc. 14, 1083–1097. 10.1175/WCAS-D-22-0026.1.

78. Chin, D.D., and Lentink, D. (2019). Birds repurpose the role of drag and lift to take off and land. Nat. Commun. 10, 5354. 10.1038/s41467-019-13347-3.

79. Swartz, S.M., Breuer, K.S., and Willis, D.J. (2008). Aeromechanics in aeroecology: flight biology in the aerosphere. Integr. Comp. Biol. 48, 85–98. 10.1093/ICB/ICN054.

80. Ortega-Jimenez, V.M., Badger, M., Wang, H., and Dudley, R. (2016). Into rude air: hummingbird flight performance in variable aerial environments. Philos. Trans. R. Soc. Lond. B Biol. Sci. 371, 20150387. 10.1098/RSTB.2015.0387.

81. del Hoyo, J. ed. (2020). All the birds of the world (Lynx Edicions, Barcelona, Spain).

82. White, R.L., and Bennett, P.M. (2015). Elevational distribution and extinction risk in birds. PLoS One 10, e0121849. 10.1371/JOURNAL.PONE.0121849.

83. Quintero, I., and Jetz, W. (2018). Global elevational diversity and diversification of birds. Nature 555, 246–250. 10.1038/nature25794.

84. Karger, D.N., Conrad, O., Böhner, J., Kawohl, T., Kreft, H., Soria-Auza, R.W., Zimmermann, N.E., Linder, H.P., and Kessler, M. (2017). Climatologies at high resolution for the earth’s land surface areas. Sci. Data 4, 170122. 10.1038/sdata.2017.122.

85. 85. BirdLife International and Handbook of the Birds of the World (2019). Bird species distribution maps of the world. Version 2019.1. http://datazone.birdlife.org/species/requestdis.

86. Jetz, W., Thomas, G.H., Joy, J.B., Hartmann, K., and Mooers, A.O. (2012). The global diversity of birds in space and time. Nature 491, 444–448. 10.1038/nature11631.

87. 87. R Core Team (2022). R: a language and environment for statistical computing. Version 4.2.2. https://www.r-project.org/.

88. Paradis, E., and Schliep, K. (2019). ape 5.0: an environment for modern phylogenetics and evolutionary analyses in R. Bioinformatics 35, 526–528. 10.1093/BIOINFORMATICS/BTY633.

89. 89. Orme, D., Freckleton, R., Thomas, G., Petzoldt, T., Fritz, S., Isaac, N., and Pearse, W. (2018). caper: comparative analyses of phylogenetics and evolution in R. Version 1.0.1. https://cran.r-project.org/package=caper.

90. Wickham, H., Averick, M., Bryan, J., Chang, W., D’, L., Mcgowan, A., François, R., Grolemund, G., Hayes, A., Henry, L., et al. (2019). Welcome to the Tidyverse. J. Open Source Softw. 4, 1686. 10.21105/JOSS.01686.

91. 91. Schnute, J.T., Boers, N., and Haigh, R. (2021). PBSmapping: mapping fisheries data and spatial analysis tools. Version 2.73.0. https://cran.r-project.org/package=PBSmapping.

92. Hijmans, R.J. (2021). raster: geographic data analysis and modeling. Version 3.4–13. https://cran.r-project.org/package=raster.

93. Ho, L. si T., and Ané, C. (2014). A linear-time algorithm for Gaussian and non-Gaussian trait evolution models. Syst. Biol. 63, 397–408. 10.1093/sysbio/syu005.

94. 94. van Buuren, S., and Groothuis-Oudshoorn, K. (2011). mice: multivariate imputation by chained equations in R. J. Stat. Softw. 45, 1–67. 10.18637/JSS.V045.I03.

95. Ives, A.R., and Li, D. (2018). rr2: An R package to calculate R^2^s for regression models. J. Open Source Softw. 3, 1028. 10.21105/joss.01028.

96. Billerman, S.M., Keeney, B.K., Rodewald, P.G., and Schulenberg, T.S. eds. (2022). Birds of the world (Cornell Lab of Ornithology, Ithaca, NY, USA). https://birdsoftheworld.org/bow/home.

97. Sibley, C.G., and Monroe, B.L. (1990). Distribution and taxonomy of birds of the world (Yale University Press, New Haven).

98. Bowler, D.E., and Benton, T.G. (2005). Causes and consequences of animal dispersal strategies: relating individual behaviour to spatial dynamics. Biol. Rev. 80, 205–225. 10.1017/S1464793104006645.

99. Freckleton, R.P. (2002). On the misuse of residuals in ecology: regression of residuals vs. multiple regression. J. Anim. Ecol. 71, 542–545. 10.1046/J.1365-2656.2002.00618.X.

100. Hackett, S.J., Kimball, R.T., Reddy, S., Bowie, R.C.K., Braun, E.L., Braun, M.J., Chojnowski, J.L., Cox, W.A., Han, K.-L., Harshman, J., et al. (2008). A phylogenomic study of birds reveals their evolutionary history. Science 320, 1763–1768. 10.1126/SCIENCE.1157704.

101. 101. Nakagawa, S., and De Villemereuil, P. (2019). A general method for simultaneously accounting for phylogenetic and species sampling uncertainty via Rubin’s rules in comparative analysis. Syst. Biol. 68, 632–641. 10.1093/sysbio/syy089.

102. Glazier, D.S. (2013). Log-transformation is useful for examining proportional relationships in allometric scaling. J. Theor. Biol. 334, 200–203. 10.1016/J.JTBI.2013.06.017.

103. Gelman, A. (2008). Scaling regression inputs by dividing by two standard deviations. Stat. Med. 27, 2865–2873. 10.1002/SIM.3107.

104. Tobalske, B.W. (2022). Aerodynamics of avian flight. Curr. Biol. 32, R1105–R1109. 10.1016/j.cub.2022.07.007.

105. Biewener, A.A. (2022). Biomechanics of avian flight. Curr. Biol. 32, R1042–R1172.

## Supplemental references

s1. Wright, N.A., Gregory, T.R., and Witt, C.C. (2014). Metabolic ‘engines’ of flight drive genome size reduction in birds. Proc R Soc Lond B Biol Sci 281, 20132780. 10.1098/RSPB.2013.2780.

s2. Sheard, C., Neate-Clegg, M.H.C., Alioravainen, N., Jones, S.E.I., Vincent, C., MacGregor, H.E.A., Bregman, T.P., Claramunt, S., and Tobias, J.A. (2020). Ecological drivers of global gradients in avian dispersal inferred from wing morphology. Nat Commun 11, 2463. 10.1038/s41467-020-16313-6.

s3. Weeks, B.C., O’Brien, B.K., Chu, J.J., Claramunt, S., Sheard, C., and Tobias, J.A. (2022). Morphological adaptations linked to flight efficiency and aerial lifestyle determine natal dispersal distance in birds. Funct Ecol 36, 1681–1689. 10.1111/1365-2435.14056.

s4. Pennycuick, C.J. (2008). Modelling the flying bird (Elsevier, Amsterdam).

s5. Tobias, J.A., Sheard, C., Pigot, A.L., Devenish, A.J.M., Yang, J., Sayol, F., Neate-Clegg, M.H.C., Alioravainen, N., Weeks, T.L., Barber, R.A., et al. (2022). AVONET: morphological, ecological and geographical data for all birds. Ecol Lett 25, 581–597. 10.1111/ele.13898.

s6. del Hoyo, J. ed. (2020). All the birds of the world (Lynx Edicions, Barcelona, Spain).

s7. White, R.L., and Bennett, P.M. (2015). Elevational distribution and extinction risk in birds. PLoS One 10, e0121849. 10.1371/JOURNAL.PONE.0121849.

s8. Quintero, I., and Jetz, W. (2018). Global elevational diversity and diversification of birds. Nature 555, 246–250. 10.1038/nature25794.

